# HNF4α isoforms regulate the circadian balance between carbohydrate and lipid metabolism in the liver

**DOI:** 10.1101/2021.02.28.433261

**Authors:** Jonathan R. Deans, Poonamjot Deol, Nina Titova, Sarah H. Radi, Linh M. Vuong, Jane R. Evans, Songqin Pan, Johannes Fahrmann, Jun Yang, Bruce D. Hammock, Oliver Fiehn, Baharan Fekry, Kristin Eckel-Mahan, Frances M. Sladek

## Abstract

Hepatocyte Nuclear Factor 4α (HNF4α), a master regulator of hepatocyte differentiation, is regulated by two promoters (P1 and P2). P1-HNF4α is the major isoform in the adult liver while P2-HNF4α is thought to be expressed only in fetal liver and liver cancer. Here, we show that P2- HNF4α is expressed at ZT9 and ZT21 in the normal adult liver and orchestrates a distinct transcriptome and metabolome via unique chromatin and protein-protein interactions. We demonstrate that while P1-HNF4α drives gluconeogenesis, P2-HNF4α drives ketogenesis and is required for elevated levels of ketone bodies in females. Exon swap mice expressing only P2- HNF4α exhibit subtle differences in circadian gene regulation and disruption of the clock increases expression of P2-HNF4α. Taken together, we propose that the highly conserved two-promoter structure of the *Hnfa* gene is an evolutionarily conserved mechanism to maintain the balance between gluconeogenesis and ketogenesis in the liver in a circadian fashion.

## Introduction

Roughly 30% of human genes contain alternative promoters and yet the functional significance of the majority of those promoters, and the transcripts they generate, is woefully understudied. One such gene is the nuclear receptor (NR) Hepatocyte Nuclear Factor 4 alpha (HNF4α), a liver-enriched transcription factor (TF) best known as a master regulator of liver-specific gene expression and mutated in Maturity Onset Diabetes of the Young 1 (MODY1) (Sladek et al., 1990) (Lu, 2016). In mouse, HNF4α is essential for fetal liver function (Battle et al., 2006) and liver knockout (KO) mice die within six weeks of birth with a fatty liver (Hayhurst et al., 2001).

The human *HNF4A* and mouse *Hnf4a* genes are highly conserved and regulated by proximal P1 and distal P2 promoters. P1 drives the expression of transcripts containing exon 1A while P2 transcripts contain exon 1D, resulting in a loss of the N-terminal activation function 1 (AF-1). In the adult liver P1 is presumed to be the only active promoter, while during fetal liver development both P1 and P2 are active (Briançon et al., 2004; Torres-Padilla et al., 2001). The first P2-HNF4α transcript cloned, HNF4α7, was from the embryonal carcinoma cell line F9 (Nakhei et al., 1998), suggesting that it might play a role in cancer as well as fetal development. P1-HNF4α is down regulated in liver cancer and acts as a tumor suppressor (Hatziapostolou et al., 2011; Ning et al., 2010; Tanaka et al., 2006; Walesky and Apte, 2015), while overexpression of P2-HNF4α is linked to poor prognosis in hepatocellular carcinoma (HCC) (Cai et al., 2017).

To address the physiological role of P2-HNF4α, we employed exon swap mice (α7HMZ), which substitute exon 1A with exon 1D in the P1 promoter and demonstrate a subtle, albeit ill-defined role for the AF-1 domain *in vivo* (Briançon and Weiss, 2006). We compared the α7HMZ adult mice (express only P2-HNF4α) to wildtype (WT) mice (express P1-HNF4α) using RNA-seq, ChIP-seq, rapid immunoprecipitation mass spectrometry of endogenous proteins (RIME), protein binding microarrays (PBMs) and metabolomics. An orchestrated, altered hepatic transcriptome in P2-HNF4α livers reveals notable differences in cytochrome P450 transcripts and subtle differences in the hepatic circadian clock, as well as an apparent feminization of male livers. The distinct P2-HNF4α transcriptome appears to be due to altered protein-protein interactions, as well as altered chromatin binding but not differences in innate DNA binding specificity. Expression of P2-HNF4α is observed at Zeitgeber time (ZT) 9 and ZT 21 in WT adult livers, and is upregulated in *Clock*-deficient mice. The P2-HNF4α hepatic metabolome is enriched in lipids and ketone bodies while mice expressing only P1-HNF4α exhibit enhanced gluconeogenesis and lack the elevated levels of ketone bodies normally found in females. Taken together, our results suggest that expression of P2-HNF4α in the liver is an evolutionarily conserved mechanism to balance carbohydrate and fatty acid metabolism during the circadian cycle.

## Results

### The P2-HNF4α transcriptome has neither a fetal nor a cancer profile

Exon swap α7HMZ mice were verified to express P2-HNF4α, but not P1-HNF4α, RNA and protein in adult liver (Figures 1A-1C). RNA-seq of adult male livers revealed a significant difference (padj ≤ 0.01) in ∼1600 genes between WT and α7HMZ, both up- (831) and downregulated (792) in α7HMZ (Figure 1D). The most downregulated (e.g., *Scnn1a*, *Cyp2c50, Rdh16f2, Ces2e*) and upregulated genes (e.g., *Rad51b, Pcp4l1, Cyp2b13, Cyp2b9*) exhibited up nearly 30-fold effects (Figure 1E). Kegg pathway analysis revealed discrete metabolic pathways in α7HMZ versus WT livers, suggesting a purposeful alteration in gene expression (Figure 1F). For example, cell adhesion molecules, drug and linoleic acid metabolism and steroid hormone biosynthesis genes were enriched in WT mice while the ribosome, oxidative phosphorylation and RNA transport and processing were enriched in α7HMZ. Several disease pathways were also enriched among the P2-specific genes, including non-alcoholic fatty liver disease (NAFLD), viral carcinogenesis, alcoholism and neurological diseases (Parkinson’s, Huntington’s, Alzheimer’s) (Figure 1F). Comparison of α7HMZ versus WT differentially expressed genes (DEGs) with HNF4α liver KO expression data (Walesky et al., 2013) revealed that ∼100 of the WT-specific genes (but none of the α7HMZ-specific genes) were downregulated in the HNF4α KO (Figure S1A), confirming P1-HNF4α predominance in the adult liver.

**Figure 1.**
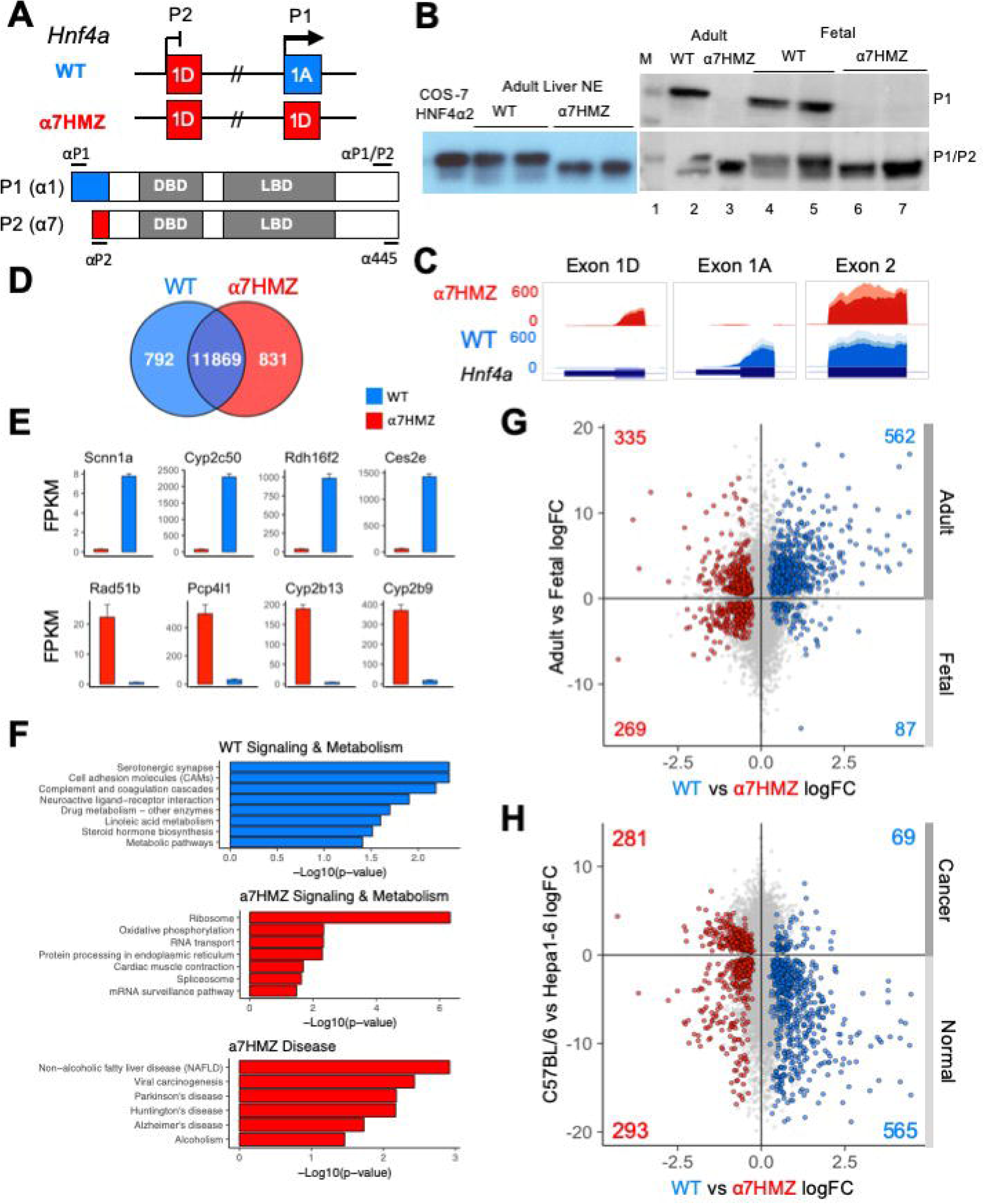
P2-HNF4α correlates with neither a fetal nor a cancer profile. (*A*) *Hnf4a* P1 and P2 promoters and first exons in WT and α7HMZ mice, protein products (P1: HNF4α1; P2: HNF4α7) and epitopes for P1-, P2-specific and P1/P2-common antibodies (Abs). DBD, DNA binding domain; LBD, ligand binding domain. (*B*) Immunoblots (IBs) of nuclear (NE) and whole cell liver extracts (WCE) with P1-specific and P1/P2-HNF4α Abs. M, molecular weight markers (top band, 54 kD). COS-7 α2, NE of cells transfected with human HNFα2. (*C*) UCSC Genome Browser view of liver RNA-seq reads mapping to *Hnf4a*. (*D*) Number of common and significantly (padj ≤ 0.01) dysregulated genes in WT and α7HMZ male liver RNA-seq (10:30 AM, ZT3.5) with baseMean ≥ 10 (n=3). (*E*) Average FPKM of most significantly up- and downregulated genes in α7HMZ livers compared to WT. All, padj <0.01. (*F*) Enriched KEGG pathways for WT- and α7HMZ-uniquely expressed genes. (*G*) RNA-seq log2 fold-change (log2FC) values between WT and α7HMZ, plotted against adult and E14.5 fetal mouse livers from ENCODE. Colored data points (total number noted in each quadrant), padj ≤ 0.01 in both datasets: blue, up in WT; red, up in α7HMZ. (*H*) As in (*G*) except plotted versus data from murine hepatoma cell line (Hepa1-6) and WT C57BL/6 livers. See Table S1 and Table S2BD for full lists of genes and Figure S1 for additional comparisons.

The α7HMZ versus WT DEGs were compared with adult versus fetal (E14.5) liver DEGs (Figure 1G). More than two thirds of the genes upregulated in WT livers were also upregulated in the adult liver, while ∼10% were enriched in fetal livers (562 versus 87, respectively). In contrast, α7HMZ-upregulated genes were more evenly split between adult and fetal liver (Figure 1G). Interestingly, alpha-fetoprotein (*Afp*) and other fetal liver genes were expressed at a lower level in α7HMZ (Figure S1B), suggesting that the α7HMZ “program” is not simply a fetal one. Furthermore, while α7HMZ mice have a significantly (p < 0.01) higher liver-to-body weight ratio than WT or α1HMZ (exon 1A swapped for exon 1D in the P2 promoter) at postnatal day 14, the reverse (α1HMZ > α7HMZ) is observed at postnatal day 21 (Figure S1C). Finally, proliferation genes *Mki67* and *Pcna* were not upregulated in α7HMZ adult livers as one might anticipate for a predominantly fetal TF (Figure S1E).

To determine whether α7HMZ livers exhibit a cancer profile, the α7HMZ versus WT DEGs were plotted against DEGs of normal C57BL/6 livers versus murine hepatoma cell line Hepa1-6 (Figure 1H). As anticipated, genes upregulated in the WT liver were preferentially expressed at higher levels in normal liver compared to liver cancer (565 versus 69, respectively). In contrast, genes more highly expressed in α7HMZ livers were not enriched in liver cancer (Figure 1H and S1D). Taken together, these results indicate that P2-HNF4α drives a specific program of gene expression in the adult liver distinct from that of P1-HNF4α that is neither completely fetal-nor cancer-like, suggesting an alternative role for P2-HNF4α.

### P2-HNF4α livers are less sensitive to the circadian clock

Since NRs are known to play an important role in regulating the circadian clock in the liver (Tahara and Shibata, 2016; Zhao et al., 2014), RNA-seq of WT and α7HMZ livers was performed at three different time points (10:30, 13:30, 20:30, equivalent to zeitgeber time (ZT) 3.5, 6.5 and 13.5, respectively). While the expression of ∼250 to 500 genes was significantly altered (padj <0.01, absolute log2FC ≥ 1) between any two of the time points in WT mice, less than half that number was altered in α7HMZ livers (Figure 2A, *top*), suggesting a reduced sensitivity of α7HMZ livers to the circadian clock. There were also more genes down-than upregulated in α7HMZ livers (Figure 2A, *bottom*), consistent with the loss of AF-1 function *in vitro* (Nakhei et al., 1998; Torres-Padilla et al., 2002). A volcano plot of α7HMZ DEGs also shows a greater number of genes increased in WT compared to α7HMZ livers at 10:30 AM (ZT 3.5), including several cytochrome P450 (*Cyp*) genes and the NR gene CAR (*Nr1i3*) (Figure 2B).

**Figure 2.**
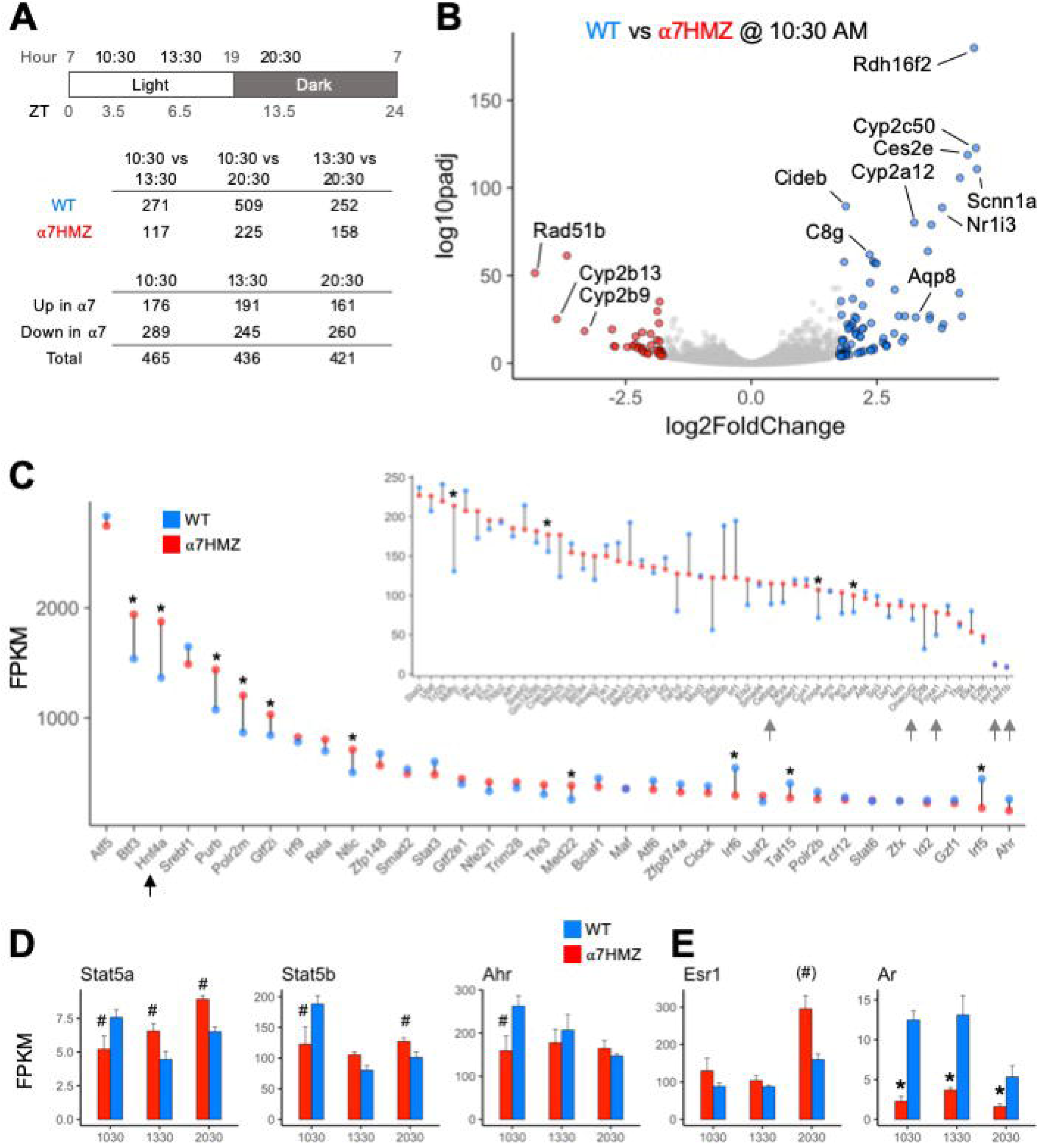
HNF4α is one of the most highly expressed TFs in the liver; P2-HNF4α promotes “feminization” of the mouse liver. (*A*) Number of genes with significant gene expression changes (padj ≤ 0.01, log2FC ≥ 1) between different time points and α7HMZ and WT in RNA-seq. (*B*) Differential gene expression at 10:30 AM. Colored spots, log2FC ≥ 1.75 (blue, WT; red, α7HMZ). (*C*) Average FPKM values for the top 85 expressed TFs in WT fed livers at 10:30 AM, sorted by WT FPKM values at 13:30. Arrows, LETFs. * padj ≤ 0.01. (*D,E*) Average FPKM of indicated genes. # padj ≤ 0.05; (#) padj = 0.055 (p=0.01); * padj ≤ 0.01. See also Figure S2.

To examine the impact of P2-HNF4α on other NR genes, we compared the FPKM values of all NR genes across all three time points. HNF4α was the most highly expressed NR in both WT and α7HMZ; the next most abundant NR, *Rxra*, was expressed at roughly 25% the level of *Hnf4a* (Figure S2A). While most NRs displayed similar circadian oscillations in WT and α7HMZ livers, there were some notable exceptions: CAR (*Nr1i3*) was highly downregulated in α7HMZ at all three time points (Figure S2A, *arrow*). Rev-Erbβ (*Nr1d2*), RORγ (*Rorc*) and PPARα (*Ppara*), all involved in the transcriptional feedback loop that drives circadian expression in the liver (Tahara and Shibata, 2016), exhibited significantly reduced expression in α7HMZ livers at one time point (Figures S2B), again suggesting a decreased responsiveness to the clock.

### HNF4α is one of the most highly expressed TFs in the liver

Consistent with the relative abundance of HNF4α protein in the adult liver (Bolotin et al., 2011; Sladek et al., 1990), HNF4α had one of the highest transcript levels of any TF, higher even than subunits of RNA polymerase II (e.g., *Polr2m*, *Polr2b*) (Figure 2C). The other liver-enriched TFs (LETFs, *Cebpa*, *Onecut2*, *Foxa1*, *Hnf1a*, *HNF1b*) had transcript levels at least 10-fold lower than *Hnf4a* (Figure 2C, *inset*, *arrows*), consistent with HNF4α being a major regulator of liver-specific gene expression. Several TFs showed statistically significant differences between WT and α7HMZ (Figure 2C, *asterisk*), including those known to play a role in sexual dimorphic gene expression (*Stat5a*, *Stat5b*, *Ahr*, *Nr0b2*) (Figure S2D and S2B) (Oshida et al., 2016; Clodfelter et al., 2007). Interestingly, *Esr1* (estrogen receptor alpha, ERα) expression was modestly upregulated in α7HMZ while *Ar* (androgen receptor, AR) was notably downregulated (Figure 2E), suggesting a potential “feminization” of the α7HMZ liver.

### P2-HNF4α dysregulates the expression of genes involved in fatty acid, steroid and xenobiotic/drug metabolism

HNF4α is a known regulator of Phase I and Phase II enzymes (Hwang-Verslues and Sladek, 2010) and has been computationally linked to sexually dimorphic and circadian expression of those genes (Hirao et al., 2011). Therefore, we examined the level of expression of all cytochrome P450 (*Cyp*) genes (Phase I) as well as glutathione S-transferases (*Gst*) and UDP glucuronosyltransferases (*Ugt*) (Phase II). While the diurnal pattern of expression was generally the same in WT and α7HMZ, the absolute level of expression was often altered (Figure 3A). For example, the expression of *Cyp2c50* and *Cyp2c54*, which encode enzymes that metabolize linoleic acid, the endogenous HNF4α ligand (Yuan et al., 2009), was much lower in α7HMZ livers (Figures 3A and 1E). Several *Ugt* genes were dysregulated and metabolomic analysis revealed a significant (padj <0.01) decrease in UDP glucuronic acid in α7HMZ livers (Figure 3A, *bottom*). Since glucose is needed to make UDP glucuronic acid, this decrease could be linked to carbohydrate metabolism (see Figure 7).

**Figure 3.**
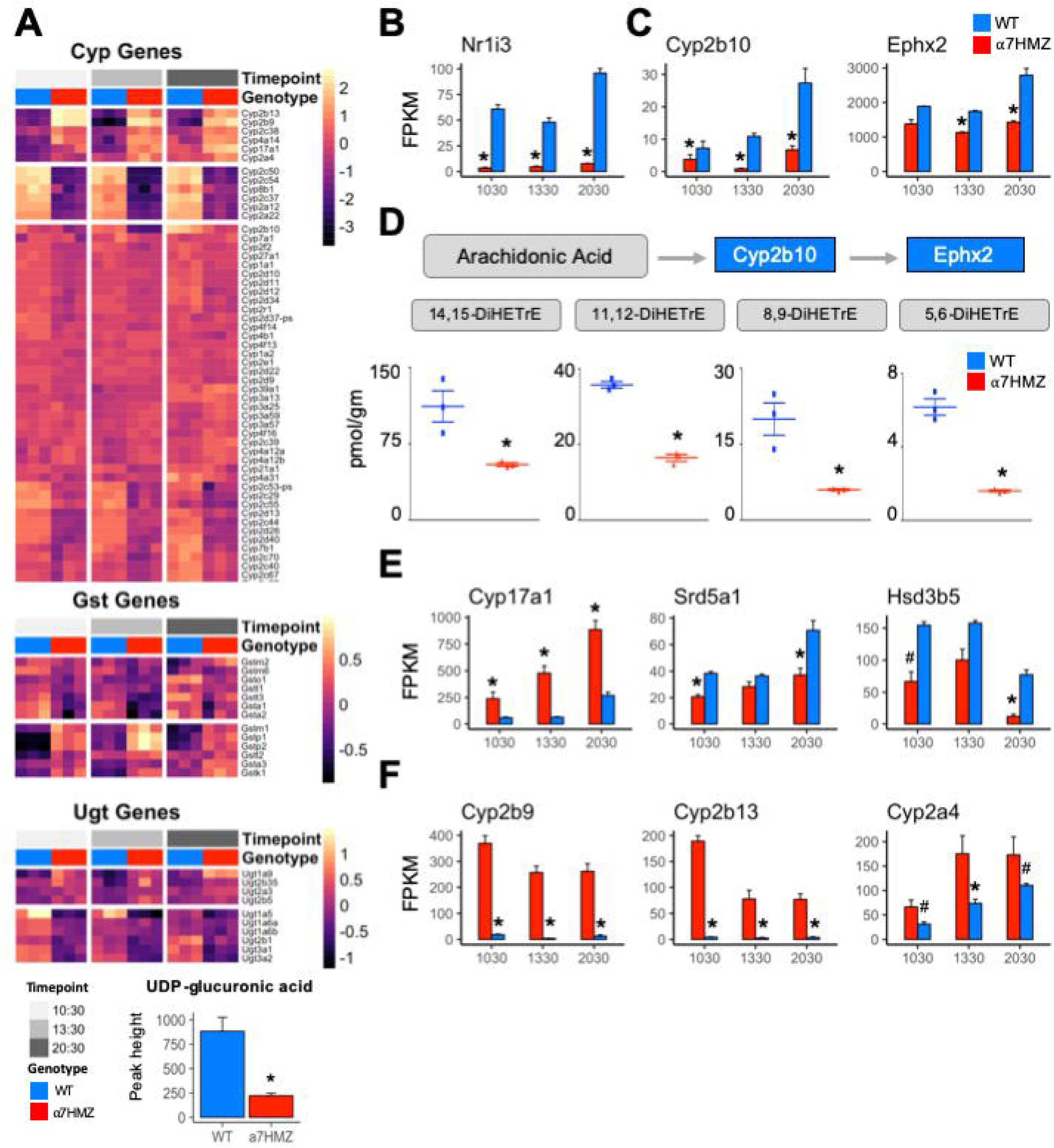
P2-HNF4α dysregulates the expression of genes involved in fatty acid, steroid and xenobiotic/drug metabolism. (*A*) Heatmap of row-normalized rlog read counts from RNA-seq for Phase I and II genes (padj ≤ 0.01) between WT and α7HMZ at any time point. *Bottom*, level of UDP-glucuronic acid in WT and α7HMZ livers, * p ≤ 0.01 by Mann-Whitney. (*B,C,E,F*) Average FPKM values. * padj ≤ 0.01 and # padj ≤ 0.05 between WT and α7HMZ at a given time point. (*D*) DiHETrE oxylipins (or DHETs, dihydroxytrienoic acid) levels in WT and α7HMZ livers (n=3, 12-13 weeks old), generated from arachidonic acid by CYP2B10 and EPHX2. * p ≤ 0.01 by Student’s T-test.

The NR CAR (*Nr1i3*) is downregulated 10- to 18-fold in α7HMZ mice (Figure 3B), as reported previously (Briançon and Weiss, 2006), and could explain some of the changes in *Cyp* gene expression observed in α7HMZ livers (Tolson and Wang, 2010). In contrast, the expression of PXR (*Nr1i2*), which is known to co-regulate many Phase I and II genes with CAR and to be upregulated by HNF4α in fetal liver (Kamiya et al., 2003; Tolson and Wang, 2010), was not altered (Figure S2C), suggesting that the primary role of P2-HNF4α in the adult liver may not be to regulate xenobiotic metabolism.

In addition to *Cyp2c50/54*, transcript levels of other fatty acid metabolic enzymes were also decreased in α7HMZ livers. *Cyp2b10* and *Ephx2* (Figure 3C), which convert arachidonic acid to oxylipins via a two-step process (Wagner et al., 2011), were significantly downregulated, as were all four DiHETrE products of arachidonic acid in the CYP2B10-EPHX2 pathway (Figure 3D), confirming a phenotypic effect on fatty acid metabolism. Changes in gene expression in the steroid metabolism pathway were also observed in α7HMZ livers with an increase in *Cyp17a1* and a decrease in *Srd5a1* and *Hsd3b5* (Figure 3E). CYP17A1 plays a predominant role in steroid hormone biosynthesis, while steroid 5-alpha-reductase (*Srd5a1*) metabolizes the conversion of testosterone into the more potent dihydrotestosterone (DHT) and 3 beta-hydroxysteroid dehydrogenase type 5 (*Hsd3b5*) is typically lower in female livers (Conforto and Waxman, 2012). Tellingly, several of the most significantly increased transcripts in α7HMZ livers, including *Cyp2b9*, *Cyp2b13 and Cyp2a4* (Figure 3F), are female-specific, have testosterone hydroxylase activity and are known to be regulated by HNF4α (Wiwi et al., 2004). Furthermore, *Ephx2* expression and activity is downregulated by estrogen (Huang and Sun, 2018), which could explain the observed decrease in DiHETrEs in α7HMZ livers. All told, these results are consistent with the “feminization” of the α7HMZ livers suggested by the increase in ERα and decrease in AR expression (Figure 2E).

### P1- and P2-HNF4α isoforms have similar but non-identical DNA binding profiles both *in vivo* and *in vitro*

To determine whether the P2-HNF4α transcriptional program is due to alterations in chromatin binding, ChIP-seq analysis was performed at 10:30 AM (ZT3.5) using an antibody (α445) that recognizes both isoforms (Figure 1A). Consistent with the high level of expression of the *Hnf4a* gene, there was a large number of HNF4α binding events in both WT and α7HMZ livers (∼40,000 peaks). While the vast majority of peaks were similar in the two sets of mice, ∼1.4 to 2.6% of the peaks were enriched for a particular isoform (WT unique: 572 peaks; α7HMZ unique: 1067 peaks) (Figure 4A). Analysis of the feature distribution of the ChIP peaks shows that both WT- and α7HMZ-unique peaks were less frequently located in the promoter region ( ≤ 2kb from +1) than the common peaks and the α7HMZ-unique peaks were enriched in intronic regions (Figure 4B).

**Figure 4.**
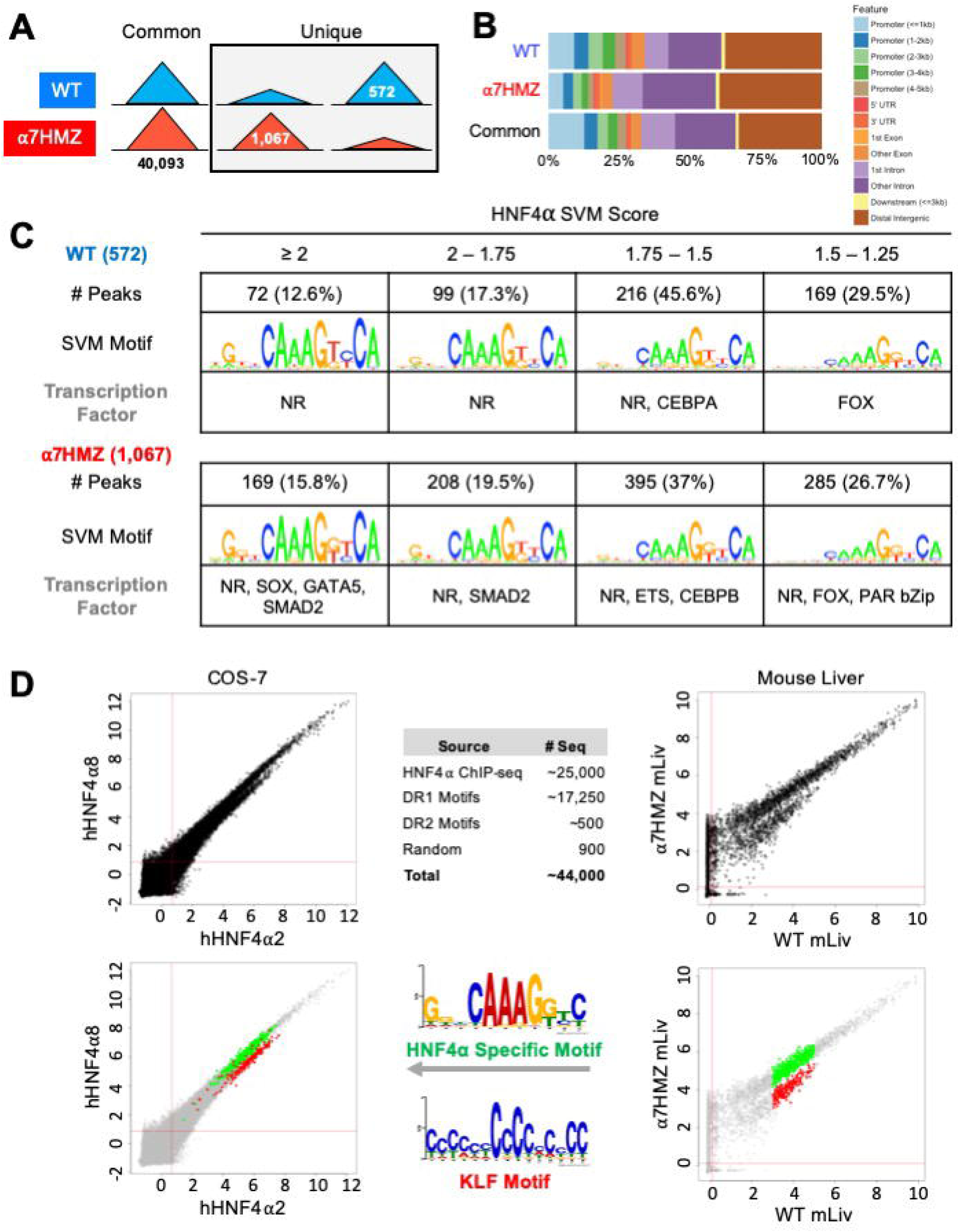
HNF4α isoforms exhibit similar but non identical chromatin binding profiles *in vivo* and *in vitro*. (*A*) Number of common, WT- and α7HMZ-unique HNF4α ChIP-seq peaks. (*B*) Feature distribution plots from ChIPseeker. (*C*) WT- and α7HMZ-unique ChIP peaks grouped by HNF4α SVM motif score. TFs corresponding to the top DNA motifs from *de novo* MEME-ChIP analysis are given. NRs, HNF4α DR1-like motif. (*D*) Log2 average binding intensities from PBMs with ectopically expressed human HNF4α8 versus HNF4α2 in COS-7 cells, and mouse liver NEs from α7HMZ versus WT. *Middle top*, test sequences used in PBM design. *Middle bottom*, PWMs for red and green spots in mouse liver scatterplot, which are mapped back onto COS-7 plots. See also Figures S3 and S4.

Motif mining showed that the most common motif in both the WT and α7HMZ unique peaks was an HNF4α motif (xxxxCAAAGTCCA). To determine whether there might be additional TFs bound in those peaks, we analyzed the DNA sequence of the uniquely bound peaks with an HNF4α-trained support vector machine (SVM) algorithm and categorized the peaks into one of four categories (>2, 2 to 1.75, 1.75 to 1.5 and 1.5 to 1.25 SVM score) based on the single highest-scoring SVM motif within the peak. All but a few peaks fell into one of these categories suggesting that the isoform-specific peaks are likely due to direct binding to the DNA. Nonetheless, *de novo* motif calling with MEME-ChIP revealed different TF motifs in some of the isoform-specific peaks. CEBPA and FOX were the only motifs significantly enriched in WT-unique peaks, but several motifs were found in α7HMZ-unique peaks, including SOX, GATA5, SMAD2, ETS, CEBPB, FOX and PAR bZIP (Figure 4C).

To investigate the innate DNA binding specificity of the HNF4α isoforms, we designed PBMs with variations on HNF4α consensus motifs (a direct repeat with a spacing of 1, DR1, AGGTCAxAGGTCA, or DR2, AGGTCAxxAGGTCA), as well as genomic sequences mined from HNF4α ChIP-seq peaks (Figure 4D, *top middle*). In total, ∼44,000 test sequences were spotted in quadruplicate on a glass slide and probed with human HNF4α2 or HNF4α8 ectopically expressed in COS-7 cells or with liver nuclear extracts (NEs) from WT and α7HMZ mice (HNF4α2/α8 have a 10-amino acid insertion in the F domain of HNF4α1/α7, respectively.) Scatter plot analysis of the PBM scores verified that the two HNF4α isoforms exhibited nearly identical DNA binding affinity and specificity across all test sequences in the COS-7 extracts (Figure 4D, *top left*). Liver NEs from WT and α7HMZ mice were also nearly identical except for a subset of sequences that differed between WT and α7HMZ (Figure 4D, *top right*). Motif analysis of the two groups of sequences, shown in green and red, revealed a preference for HNF4α in WT livers for GC-rich sequences recognized by SP1/KLF proteins (Figure 4D, *bottom right, middle*). A similar, albeit less pronounced, preference was noted in the COS-7 extracts (Figure 4D, *bottom left*). Consistently, HNF4α1 has been found to interact with SP1 both on and off chromatin, an interaction that involves the N-terminal domain of HNF4α1 (Hwang-Verslues and Sladek, 2008; Kardassis et al., 2002; Takahashi et al., 2002).

### HNF4α isoforms have unique interactomes

To assess the contribution of differential chromatin binding to changes in gene expression, we cross-referenced the ChIP-seq and RNA-seq datasets at the 10:30 AM time point and found that ∼22% of WT-specific (62 out of 294) and α7HMZ-specific genes (41 out of 181) have one or more unique ChIP peaks within 50 kb of the transcription start site (TSS, +1) (Figure 5A). WT-specific genes matching these criteria include *Nr1i3*, *Cyp2c50*, *Cyp2c54*, *Rarres1*, *Fmn1*, *Cdhr5*, and *Camk1d*, while α7HMZ-specific genes include *Cyp2b9*, *Fgfr1*, *Wnk4*, *Cyp4a14*, *Ppl*, *Vnn1*, *Acot1*, and *Cyp17a1* (Table S2E and S2F). Many of the most dysregulated genes contained differentially bound peaks within ∼5 kb of +1 -- *Nr1i3* (CAR), *Apoa4, Cyp2c50, Cyp2c54, Cyp2b9, Cyp4a14, Acot1, Cyp17a1,Ucp2, Cyp2d26* and *Treh* (Figures 5B, 5C and S4AB). While the differential peaks were typically not the only nor the largest peak in the gene, they could reflect rapid cycling on and off the DNA with functional consequences.

**Figure 5.**
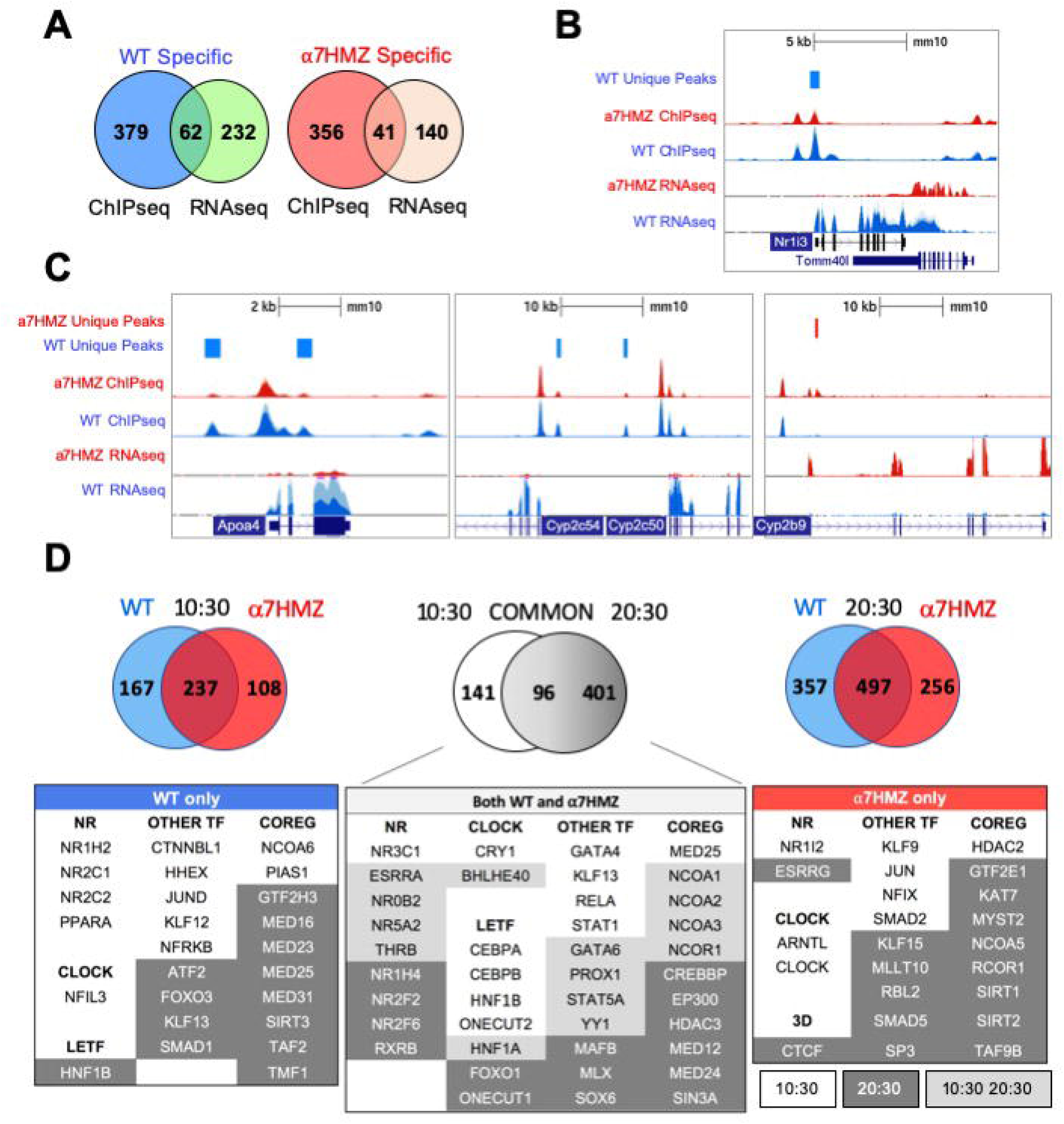
HNF4α isoforms have unique protein-protein interactions. (*A*) Number of genes with one or more WT- or α7HMZ-unique ChIP-peaks within a 50-kb of +1 of differentially expressed genes in WT and α7HMZ livers (padj ≤ 0.01). (*B,C*) UCSC Genome Browser view of dysregulated genes with a unique ChIP-signal and RNA-seq from 10:30 AM. Axes for WT and α7HMZ signals are set to the same scale but may differ between genes. (*D*) *Top*, number of proteins bound to HNF4α in WT vs. α7HMZ livers at 10:30 (ZT3.5 *left*), at 20:30 (ZT13.5 *right*) and in both WT and α7HMZ at 10:30 vs. 20:30 (*middle*). *Bottom*, select proteins involved in transcription regulation bound only in WT, α7HMZ or both genotypes. See Table S3 for all interacting proteins and Figure S4D.

Since the majority of dysregulated genes had no nearby HNF4α isoform-specific ChIP peak, we examined HNF4α protein-protein interactions in WT and α7HMZ livers by RIME at 10:30 (ZT3.5) and 20:30 (ZT13.5). Both time points yielded a considerable number of interacting proteins at least eight-fold above the background, including many proteins that bound a single isoform (10:30: 167 WT-specific, 108 α7HMZ-specific; 20:30: 357 WT-specific, 256 α7HMZ-specific) (Figure 5D, *top*). There was considerable overlap between the common groups for the two time points (96 proteins bound HNF4α in both WT and α7HMZ livers at both time points), underscoring the robustness of the method. There were also many proteins that bound both isoforms but only at a single time point (10:30: 141; 20:30: 401). Notably, core circadian regulator CRY1 bound HNF4α in both WT and α7HMZ livers but only at 10:30 AM (Figure 5D, *bottom*). In contrast, BHLHE40 bound both isoforms at both 10:30 and 20:30, while NFIL3 uniquely bound in WT livers and ARNTL (BMAL1) and CLOCK in α7HMZ but only at 10:30 (Figure 5D, *bottom*). These findings are consistent with recent reports of HNF4α interacting with the clock machinery and playing a role in maintaining circadian oscillations in the liver (Qu et al., 2018).

Several LETFs interacted with both isoforms but only at one time point (10:30 only: CEBPA, CEBPB, HNF1B, ONECUT2; 20:30 only: FOXO1, ONECUT1); only HNF1A interacted at both time points (Figure 5D, *bottom, middle*). Many NRs interacted with HNF4α in both WT and α7HMZ livers, including NR3C1(GR) and NR0B2 (SHP), both of which have been shown previously to functionally interact with HNF4α, further validating the RIME results (Hall et al., 1995; Lee et al., 2000). There were also isoform-specific interactions, mostly with WT at 10:30 (NR1H2, NR2C1, NR2C2, PPARA). Interestingly, xenobiotic receptor PXR (NR1I2) interacted with HNF4α but only in α7HMZ livers at 10:30 (Figure 5D, *bottom left* and *right*). While the expression of *Nr1i2* was not changed in α7HMZ livers (Figure S2C), an environmental estrogen that activates PXR has been shown to increase the expression of two female-specific Cyp genes (*Cyp2b9* and *Cyp2a4*) in male mice: both are highly upregulated in α7HMZ livers and bound by HNF4α (Figure 5C and Table S2E) (Hernandez et al., 2006). HNF4α in α7HMZ livers also uniquely interacted with ESRRG (ERRγ, *Nr3b3*) but only at 20:30: ERRs play important roles in mitochondrial biogenesis and function, including fatty acid oxidation (Hock and Kralli, 2009). Interestingly, ERR DNA binding motifs were found in α7HMZ ChIP peaks but not WT (Figure S3B).

There were many other TFs and co-regulators that interacted with a single HNF4α isoform, often in a circadian fashion (Figure 5D, *bottom*), which could explain the observed differential gene expression between WT and α7HMZ. Several of these proteins were previously confirmed by more conventional means (Maeda et al., 2002; Ruse et al., 2002; Sladek et al., 1999; Torres-Padilla et al., 2002). Finally, there were several signaling molecules that interacted uniquely with the isoforms and at distinct time points -- one or more of these could also contribute to isoform-specific gene expression, independent of ChIP peaks (Figure S4D).

### HNF4α isoforms differentially impact circadian gene expression

Interactions with circadian TFs suggest that HNF4α may play a role in the hepatic clock. Analysis of all DEGs between any two time points (10:30, 13:30 or 20:30; padj<0.01 and log2FC>2) for either WT or α7HMZ yielded 53 genes, including commonly known circadian genes (*Cry1*, *Rorc*, *Dbp*, *Bhlhe41*, *Usp2*, *Per2*, *Per3*, *Arntl*, and *Nr1d1*) as well as many metabolism-related genes (*Fmo3*, *Lpl*, *Car3*, *Corin*, *Npas2*, *Hmgcs1*, *Mme*, *Slc45a3*, *Hsd3b4*, *Hsd3b5*, *Slc10a2*) (Figure 6A *top*). While most of these circadian-regulated genes showed the same general profile in WT and α7HMZ livers, there were some differences in the magnitude of the circadian effect between the genotypes. For example, *Fmo3*, a drug metabolizing gene whose expression varies greatly between individuals, had a much higher expression in α7HMZ livers at 20:30. In contrast, *Aqp8,* a water channel protein important for mitochondrial respiratory function (Ikaga et al., 2015), had much lower levels of expression in α7HMZ at all time points (Figure 6A, *arrows*).

**Figure 6.**
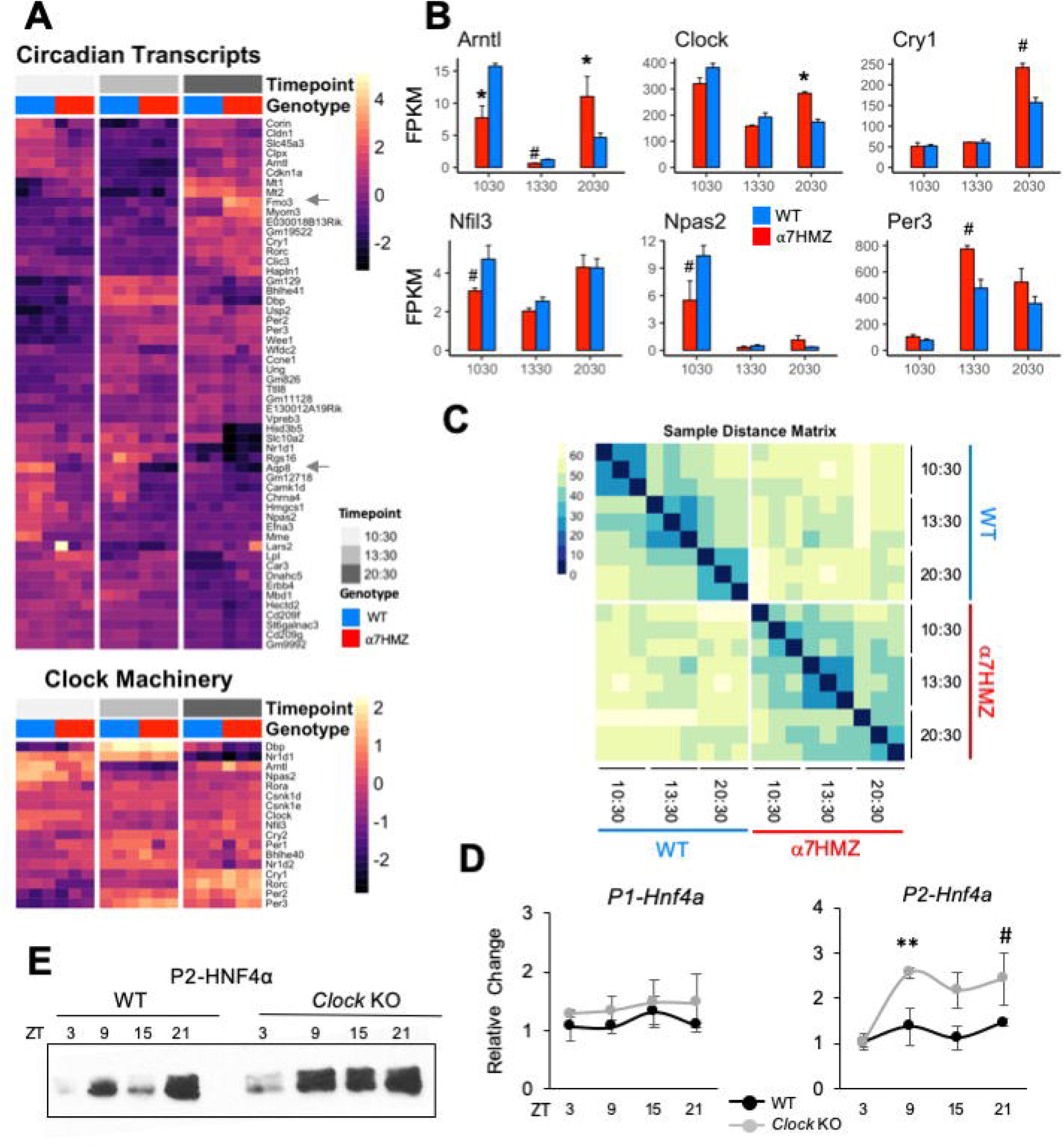
HNF4α isoforms differentially impact diurnal gene expression. (*A*) Heatmap of rlog read counts for all circadian-regulated and core clock genes with padj ≤ 0.01 and log2FC ≥ 2 between any pair of time points. (*B*) Average FPKM of select circadian clock genes. # padj ≤ 0.05; * padj ≤ 0.01. (*C*) Sample distance matrix for each RNA-seq replicate, calculated across the full transcriptome. The darker the color, the higher the degree of similarity. (*D*) Relative fold increase in P1- and P2-HNF4α in livers of WT or littermate *Clock* KO by qRT-PCR at indicated ZT. Two-way ANOVA, Sidak’s multiple comparisons test, # p ≤ 0.05; ** p ≤ 0.005. Error bars, SEM (n=3-4). (*E*) IB of diurnal expression of P2-HNF4α protein in WCE from WT and *Clock* KO livers using P2-specific antibodies. See also Figure S5.

While the majority of the clock machinery maintained cyclic expression in both genotypes (Figure 6A *bottom*), there were significant differences in expression between WT and α7HMZ in core clock components *Arntl*, *Clock*, *Cry1*, *Nfil3*, *Npas2* and *Per3* at one or more time points (Figure 6B), as well as *Rorc* and *Ppara* (Figure S2B). The fact that other core components of the clock machinery did not show differences between the two genotypes (e.g., *Per1*, *Per2*, *Rora, Bhlhe40*) (Figure S5A) suggests that the effect of P2-HNF4α on the clock is a specific one.

A sample distance matrix further confirmed a subtle yet real effect of P2-HNF4α on the hepatic clock. While the WT replicates at a given time point are much more similar to each other than they are to other time points, α7HMZ replicates show strong self-identity only in the 13:30 samples (Figure 6C). This is despite the fact that a principal component analysis (PCA) showed a good separation and categorization of each sample group (Figure S7A).

### P2-HNF4α is expressed at discrete times in the normal adult liver

While expression of P2-HNF4α protein in the normal adult liver has not been previously reported, this could be due to the time of day that livers are typically harvested (before midday). Since the current results show links between P2-HNF4α in α7HMZ livers and the circadian clock, we harvested livers from WT mice at four time points (ZT3, ZT9, ZT15, ZT21) and looked for P2-HNF4α mRNA by qPCR and protein by immunoblot (IB). The results show expression of P2-HNF4α at ZT9 (4 pm) and ZT21 (4 AM) and a further increase in *Clock* KO livers. In contrast, P1-HNF4α did not show a significant circadian effect in either WT or CLOCK KO mice (Figures 6D, 6E, S5B, S5C).

### Metabolomic profiling indicates a role for P2-HNF4α in ketogenesis

Since metabolism is tightly linked to the clock in the liver (Eckel-Mahan et al., 2012, 2013; Ribas-Latre and Eckel-Mahan, 2016), we performed metabolomic analysis of primary metabolites and complex lipids on WT and α7HMZ livers at 10:30. Approximately one-quarter of the primary metabolites (100 out of 359 total) were significantly down-regulated (p <0.05) in α7HMZ livers (Figure 7A, up in WT). Metabolite Set Enrichment Analysis showed that the top four enriched categories in WT are involved in carbohydrate metabolism and protein biosynthesis (Figure S6A). Glucose and pyruvate were both significantly down in α7HMZ livers (Figure 7B), as was PEPCK (*Pck1*), an important enzyme in gluconeogenesis (Figure S6B). Genes in pathways downstream of pyruvate were also significantly decreased in α7HMZ, including lactate dehydrogenase (*Ldha*, *Ldhd*), pyruvate carboxylase (*Pcx*) and citrate synthase (*Cs*) in the Kreb’s cycle (Figure S6C), as was citric acid, a key intermediate in the cycle (Figure 7B). Kreb’s intermediates oxalic and succinic acid were also reduced although they did not reach significance (Figure S6D). In contrast, genes involved in the formation of ketone bodies were upregulated in α7HMZ (*Hmgcs2*, *Hmgcl*) (Figure S6E), as was the ketone body β-hydroxybutyric acid (Figure 7B), as previously reported (Briançon and Weiss, 2006). Levels of hundreds of complex lipids were altered (up or down) in α7HMZ livers, including a notable increase in total triglycerides, diacylglycerides and acylcarnitines in the α7HMZ liver and a decrease in phospholipid species (Figures 7AC and S6F).

**Figure 7.**
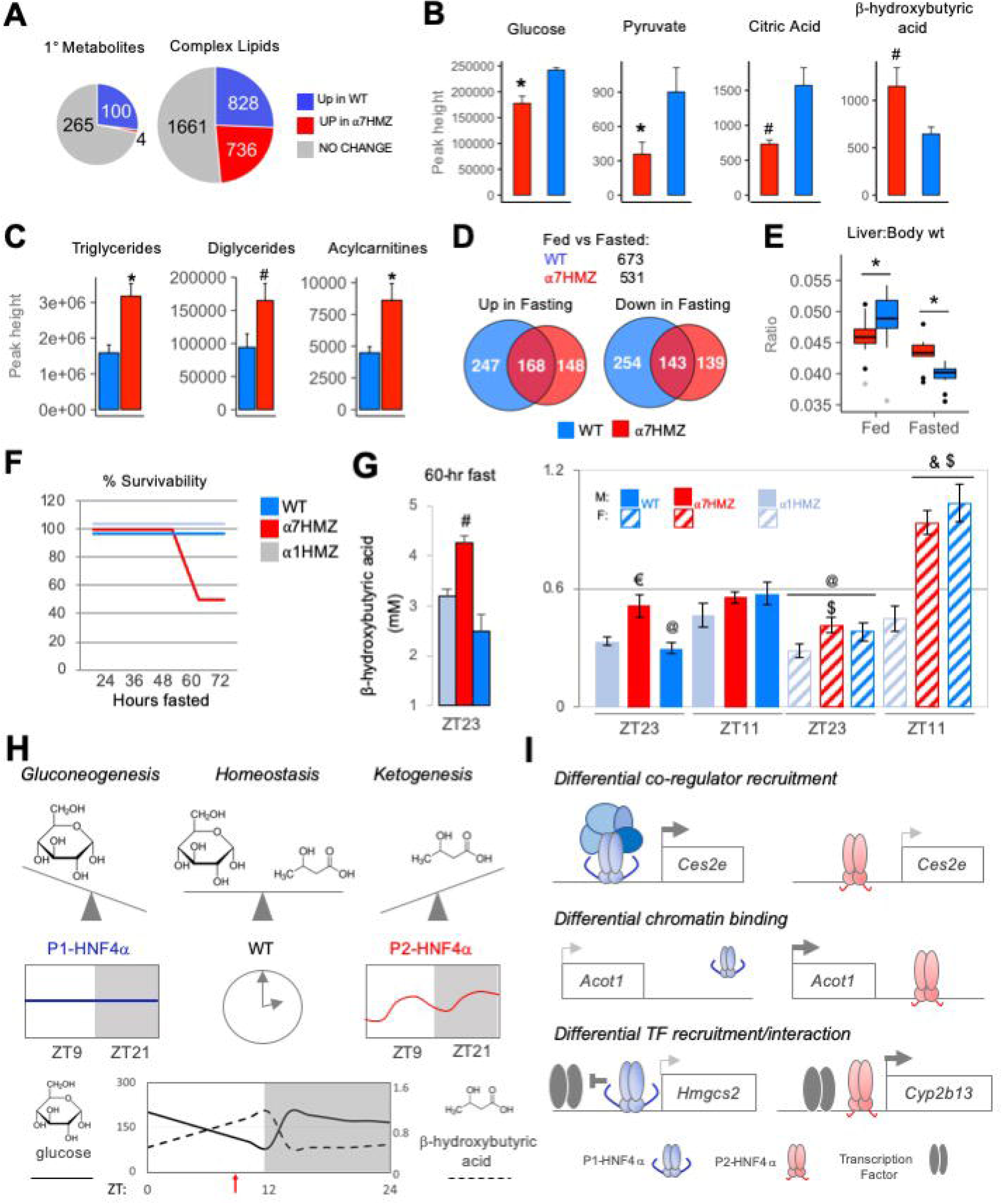
P1-HNF4α drives gluconeogenesis while P2-HNF4α drives ketogenesis; both are required for homeostasis in carbohydrate and lipid metabolism. (*A*) Number of primary metabolites and complex lipids in WT and α7HMZ livers of fed mice at 10:30 AM (n=8). Uniqueness identified by Mann-Whitney U-test (p ≤ 0.05). (*B*) Primary metabolites related to Krebs cycle and ketogenesis (a single mouse outlier was omitted for both the WT and α7HMZ datasets for a final n=7) and known complex lipids (n=8) (*C*). Student’s T-test: # p ≤ 0.05; * p ≤ 0.01. (*D*) Number of genes dysregulated (padj ≤ 0.01) in fed vs. fasted (12 hr) WT and α7HMZ livers (10:30 AM). (*E*) Liver-to-body weight ratios in fed and fasted WT and α7HMZ mice (n = 9-14). * p ≤ 0.01 by Student’s T-test, excluding outliers (gray dots). (*F*) Percent survival of WT, α1HMZ and α7HMZ male mice (∼20 wks) during a prolonged fast (n=5-6). (*G*) β-hydroxybutyric acid levels in blood of WT, α7HMZ and α1HMZ during a 60-hr fast (*left*) (n=3-5) and at ZT23 and ZT11 after 1 week of restricted feeding (food removed between ZT0 and ZT12) (*right*) (n = 6-8). Student’s T-test: # p<0.05 vs. other two genotypes; $ p<0.05 vs. α1HMZ females at the same time point; & p<0.05 vs. males of same genotype at ZT11; @ p<0.05 vs. females of the same genotype at ZT11; € p<0.05 vs. WT and α1HMZ males at ZT23. (*H,I*) Models discussed in text. See also Figures S6 and S7.

Since ketone bodies are elevated upon fasting, we performed RNA-seq on livers from 12- hr fasted WT and α7HMZ mice. The transcriptomes for both WT and α7HMZ fasted livers were quite distinct from the fed time points as well as from each other (Figures S7A and S7B): WT mice had more genes altered upon fasting (673 versus 531 in α7HMZ) as well as more WT-specific genes either up- or downregulated (Figure 7D). Liver-to-body weight ratios were significantly lower in α7HMZ versus WT fed mice; in contrast, in fasted livers the ratio was lower in WT (Figure 7E). Since WT mammals are known to store fat in their liver during periods of fasting and since fasted α7HMZ livers accumulate more fat than WT mice (Figure 7C)(Briançon and Weiss, 2006), these results suggested that P2-HNF4α might promote a “fasting-response” program, consistent with the expression of P2-HNF4α at ZT9, near the end of the daily fasting period.

When the mice were subjected to a prolonged fast, unexpectedly, 50% of the α7HMZ mice died after ∼60 hrs; in contrast, α1HMZ and WT mice survived a full 72 hrs without food (Figure 7F). Mortality was not due to hypoglycemia as blood glucose levels did not drop below 65 mg/dL; in fact, they increased after 48 hrs of fasting at ZT11, especially in α1HMZ (Figure S7C). In contrast, circulating ketone bodies were highly elevated in the α7HMZ mice that survived the 60-hr fast (4.25 mM) (Figure 7G), and suggested that the α7HMZ mice undergoing a prolonged fast died of ketoacidosis.

Since the α7HMZ transcriptome showed signs of “feminization” and since females tend to have higher levels of ketone bodies than males (Halkes et al., 2003; Marinou et al., 2011), we examined whether the elevated levels of ketone bodies in WT females is due to the ability to express P2-HNF4α. As anticipated, in WT mice ketone bodies were higher near the end of the daily fast (ZT11, 7 PM) than at the end of the feeding period (ZT23, 7 AM), in both males and females (Figure 7G). In contrast, α7HMZ males had ketone bodies at ZT23 nearly as high as at ZT11. Importantly, α7HMZ and WT females had much higher levels of ketone bodies at ZT11 than their male counterparts, whereas the α1HMZ females had levels similar to α1HMZ males and much lower than either WT or α7HMZ females (Figures 7G). This suggests that P2-HNF4α is required for the elevated levels of ketone bodies in females.

## Discussion

While many mammalian genes have multiple promoters that drive expression of proteins with alternative N-termini, the physiological relevance of those different isoforms is seldom known. Using exon-swap mice and omics approaches, we show for the first time that the alternative isoform of the *Hnf4a* gene (P2-HNF4α), previously thought to be expressed only in fetal liver and liver cancer, plays an important metabolic role in the adult liver and is implicated in both the circadian clock and sex-specific gene expression.

### Both P1- and P2-HNF4α are required for metabolic homeostasis in males and females

The “P2-HNF4α program” is characterized by a decrease in carbohydrate metabolism and an increase in hepatic fat storage and utilization, as well as ketogenesis, which typically occur during periods of fasting (Puchalska and Crawford, 2017). Altered expression of genes involved in fatty acid oxidation or oxidative phosphorylation in the mitochondria are consistent with a shift from carbohydrates to fatty acids as an energy source (e.g., *Hmgcs2*, *Acot1*, *Ucp2*, Figure 1F and Table S2G). In contrast, P1-HNF4α drives gluconeogenesis and is required to temper the P2-HNF4α response to avoid ketoacidosis. Only WT mice, which express both HNF4α isoforms in the liver, achieve homeostatic balance between carbohydrate and lipid metabolism (Figure 7H). This balance is achieved on a daily basis by upregulating P2-HNF4α at the end of the fasting period (∼ZT9) (Figure 6), resulting in the well characterized elevation of ketone bodies right before feeding (Chavan et al., 2016). Intriguingly, P2-HNF4α is also required for the elevated levels of ketone bodies in female mice (Figure 7G), consistent with a “feminization” of the α7HMZ livers and previous results showing that KO of HNF4α in the adult liver leads to a loss of male-specific genes and an increase of female-specific genes (Holloway et al., 2008). Finally, P2- but not P1-HNF4α interacts with the NAD-dependent deacetylase SIRT1, which is activated upon fasting and is associated with fatty acid oxidation, ketogenesis and fatty liver (Figure 5D) (Nassir and Ibdah, 2016).

### Multiple mechanisms are responsible for HNF4α isoform-specific gene regulation

Our results indicate that P2-HNF4α drives its unique transcriptional program via multiple mechanisms. Differential recruitment of co-regulators to target gene promoters could explain altered expression of genes such as *Ces2e*, which encodes carboxylesterase 2, an enzyme that hydrolyzes triacylglycerols (Figure 7I *top* and Figure 5D). Both P1- and P2-HNF4α bind the *Ces2e* promoter in a similar fashion but *Ces2e* is expressed at much higher levels in α7HMZ livers compared to WT, which could explain the elevated levels of triglycerides in α7HMZ livers (Figures 1E, S4C, 7C). A second potential mechanism is differential binding to regulatory regions (Figure 7I *bottom*). An enriched ChIP-seq peak in α7HMZ livers, for example, could explain the upregulation of a key enzyme in β-oxidation of fatty acids, *Acot1* (Figures 7I *middle* and S4A). A third mechanism involves differential recruitment and/or interaction of TFs with a given HNF4α isoform (Figure 7I *bottom*). For example, PPARα is known to be a major player in ketogenesis, activating the expression of the mitochondrial enzyme HMGCS2 which catalyzes the first step in ketogenesis (Puchalska and Crawford, 2017). P1-HNF4α has been shown to decrease *Hmgcs2* expression by repressing PPARα-dependent activation (Rodríguez et al., 1998). This repression could be facilitated by a unique protein-protein interaction between P1- HNF4α and PPARα (Figure 5D). In contrast, in α7HMZ livers *Hmgcs2* expression is elevated even though PPARα expression is somewhat reduced and HNF4α ChIP-seq peaks in α7HMZ livers are similar to those in WT (Figures S2B, S4C, S6E). Similarly, specific interactions between P2-HNF4α and TFs involved in sex-specific gene expression (e.g., NR1I2, SP1 family/KLF) could contribute to increased expression of female-specific genes such as *Cyp2b13* (Figures 3F, 5D, S4C) (Hernandez et al., 2006). Additional mechanisms driving the P2-HNF4α program include differential interaction with signaling molecules, altered expression of other TFs, such as those that play a role in sex-specific gene expression (Stat5b, Stat5a, estrogen and androgen receptors) (Hirao et al., 2011; Oshida et al., 2016), and elevated levels of ketone bodies which can impact histone deacetylase activity, as well as the circadian clock (Tognini et al., 2017) (Figures 2, S4D, 7G).

### Physiological and pathological triggers of the P2-HNF4α program

There are now three known physiological conditions in which P2-HNF4α is expressed in the liver -- fetal liver and ZT9 and ZT21 in adult liver (Figure 6). Increased expression of P2- HNF4α expression right before birth (E17.5), followed by a sharp decline after birth (Briançon et al., 2004; Torres-Padilla et al., 2001), could explain why the α7HMZ transcriptome is not more similar to that of the E14.5 fetal liver: rather than promoting early liver development, the role of P2-HNF4α appears to be a metabolic one, perhaps preparing the fetus to survive the birthing process and immediate postnatal period by increasing fat in the liver. The subsequent decrease in P2-HNF4α expression after birth could be mediated by GR which is induced by stress hormones released during labor (Rando et al., 2016): GR preferentially increases the expression of P1- HNF4α (Bailly et al., 2009; Nakhei et al., 1998) which would in turn repress the P2 promoter (Briançon et al., 2004).

Factors responsible for increased expression of P2-HNF4α at ZT9 have not been identified, but its expression seems to be required for the increased the level of ketogenic genes and ketone bodies in response to the daily fast (Figures S6E and 7BG). The role of P2-HNF4α at ZT21 is more difficult to explain as ketone bodies are low at that time (Figure 7G) (Chavan et al., 2016). Total protein synthesis is increased at ∼ZT22 (Robles et al., 2014), as well as both P2- and P1-HNF4α-specific targets (Figure S7D), so expression of P2- (and P1-)HNF4α at ZT21 might be the result of a global effect on protein synthesis.

In addition to physiological triggers, there are now four pathological conditions in which P2-HNF4α is known to be elevated in the adult liver -- cancer (Tanaka et al., 2006), high fat diet (Fekry et al., 2018), disrupted clock (Figure 6DE) and alcoholic hepatitis (Argemi et al., 2019). In terms of cancer, our results indicate that P2-HNF4α is not oncogenic per se -- the P2-HNF4α transcriptome shows only a partial overlap with HCC, key proliferation markers (Ki67 and PCNA) are not upregulated in α7HMZ livers, there is no evidence of hepatomegaly (Figures 1H, S1DE, 7E) and no increase in spontaneous, macroscopic tumors have been observed in α7HMZ livers, even in older mice (unpublished observation). While HCC patients with increased P2- HNF4α have a poor prognosis (Cai et al., 2017), rather than acting as an oncogene *per se*, P2- HNF4α may be upregulated simply due to a decrease in the expression of the tumor suppressor P1-HNF4α (Bailly et al., 2009; Nakhei et al., 1998) and inadvertently promote liver cancer progression via metabolic effects. For example, acylcarnitines are elevated in α7HMZ livers (Figures 7C, S6F) and have been identified as potential diagnostic and prognostic biomarkers for HCC (Lu et al., 2016; Yaligar et al., 2016). Several matrix metalloproteinases (*Mmp14*, *Mmp15*, *Mmp19*), which are linked to poor prognosis of liver or colorectal cancer patients (Chen et al., 2011, 2019; Zheng et al., 2019), are also upregulated by P2-HNF4α (Table S1). Finally, dysregulation of genes involved in drug metabolism could also impact treatment of liver cancer (Figure 3).

The second condition that leads to expression of P2-HNF4α in the adult liver -- high fat diet -- could be related to both cancer and the third condition, disrupted clock. We recently reported that P2-HNF4α expression is increased in the livers of mice fed a high fat diet and that the circadian regulator BMAL1 represses P2-HNF4α expression in HCC (Fekry et al., 2018). Consistently, P2- but not P1-HNF4α interacts with BMAL1 (ARNTL) and CLOCK and the *Clock* KO increases P2- but not P1-HNF4α expression (Figures 5D and 6DE). Dysregulation of the clock, such as during jet lag, could potentially contribute to liver cancer by upregulating P2- HNF4α (Figure 6DE) (Kettner et al., 2016).

The fourth pathological condition where P2-HNF4α is expressed in the liver -- human alcoholic steatohepatitis (Argemi et al., 2019) -- is consistent with increased fat in α7HMZ livers and an enrichment of genes associated with alcoholism in α7HMZ mice (Figures 1F, 7C). The TGFβ pathway is implicated in P2-HNF4α expression under this scenario; SMAD binding motifs were found in α7HMZ ChIP-seq peaks but not WT peaks (Figure S3B).

In summary, our results with the exon swap mice reveal important functional differences between the HNF4α isoforms and suggest a new role for P2-HNF4α in the liver. Future studies are required to identify conditions that cause the switch between the isoforms in a wildtype liver and to further explore the impact of that switch on liver physiology and disease.

## Materials & Methods

(see Supplemental Methods for additional methods and details)

### Animal models

Young adult (16 to 20 weeks) male WT and α7HMZ mice in a mixed 129/Sv plus C57BL/6 background (Briançon and Weiss, 2006) were fed a standard lab chow (LabDiet, #5001, St. Louis, MO) and used for RNA-seq, CHIP-seq, RIME analysis (all samples from the same set of mice), and oxylipin analysis. The α7HMZ male mice used for primary metabolite and complex lipid metabolomic analysis were backcrossed to C57BL/6N for 10+ generations and used with C57BL/6N WT controls (n=8, 35 weeks of age). α7HMZ and α1HMZ (backcrossed 10+ generations into C57BL/6N) were compared to C57BL/6N (WT) controls for newborn liver analysis (mixed-sex) and glucose/ketone body analysis (males and females; ∼16 to 20 weeks of age). *Clock*-deficient (*Clock* KO) male mice were provided by Dr. David Weaver (Debruyne et al., 2006). All mice were fed *ad libitum* and kept in 12-hr light/dark conditions, unless indicated otherwise, and euthanized by CO_2_ asphyxiation followed by tissue harvest at the indicated time points. Care and treatment of the animals were in strict accordance with guidelines from the Institutional Animal Care and Use Committee at the University of California, Riverside, or the McGovern Medical School, UT Health.

### Expression profiling (RNA-seq) and analysis

Next generation sequencing of RNA (RNA-seq) was carried out as previously described (Vuong et al., 2015). WT and α7HMZ male mice were sacrificed (n=3, aged 16-18 weeks) at the indicated time points -- 10:30, 13:30, 20:30 (ZT 3.5, ZT 6.5, and ZT 13.5, respectively) -- within a 30-min interval. Fasted mice had food removed from 22:30 (ZT15:30) to 10:30 AM (ZT3.5) the following day (12 hr). Libraries were submitted for 75-bp single-end sequencing with Illumina NextSeq 500 at the UCR IIGB Genomics Core. A total of 24 libraries (3 fed time points, 1 fasted time point, 2 genotypes each, 3 replicates) were multiplexed and sequenced in two separate runs, each of which yielded ∼600 M reads, averaging ∼50 M reads per sample.

Reads were aligned to the mouse reference genome (mm10) with TopHat v2.1.1 using default parameters except for allowing only 1 unique alignment for a given read. Raw read counts were calculated at the gene level for each sample using HTSeq v0.6.1. Library normalization was performed with EDASeq (Risso et al., 2011); within-lane normalization on GC content was performed with the LOESS method and between-lane normalization was performed with non-linear full quantile method. Normalization factors from EDASeq were used for differential expression analysis with DESeq2. Normalized read counts, FPKM (fragments per kilobase per million), and rlog (regularized log transformation) results were generated for downstream analysis.

### Chromatin Immunoprecipitation Sequencing (ChIP-seq) and SVM analysis

ChIP-seq of isolated liver cells from WT and α7HMZ males (n=3, aged 16-18 weeks) was performed as previously described (Vuong et al., 2015) using 4.2 µg of affinity-purified anti- HNF4α (α445) (Sladek et al., 1990) or rabbit IgG control (Santa Cruz, cat#sc-2027). Libraries were submitted for 50-bp single end sequencing by Illumina HiSEQ 2500 at the UCR IIGB Genomics core. Reads were aligned to the mouse reference genome (mm10) with Bowtie2. Peaks were called with MACS2 for individual samples, as well as a pooled peak dataset using the SPMR (signal per million reads) parameter. Aligned reads and MACS2 peak-sets were analyzed with DiffBind (Stark and Brown, 2011), with DESeq2 and library size equal to total aligned reads to identify common and uniquely bound regions of the genome. Default parameters were used unless noted otherwise. ChIP-seq peaks were called with MACS2 and then filtered on -log10(p-value) ≥ 10, to approach six-fold enrichment above control. Differentially bound peaks were identified using DiffBind with MACS2 output. Curated peak lists were generated by filtering all results on peaks with “concentration” ≥ 5; defined by DiffBind as the “mean (log) reads across all samples” in contrast. The kernel-based SVM was trained as previously described using results from independent HNF4α PBM experiments (Bolotin et al., 2010).

### Protein Binding Microarrays (PBM)

Protein binding microarrays (PBMs) were carried out as previously described (Bolotin et al., 2010). Nuclear extracts (NE) were prepared from COS-7 cells transiently transfected via CaPO_4_ with HNF4α expression vectors for human HNF4α2 (NM_00457) and HNF4α8 (NM_175914) essentially as previously described (Jiang et al., 1995). Liver NE from WT and α7HMZ mice were prepared as previously described (Yuan et al., 2009).A custom-designed array was ordered from Agilent (SurePrint G3 Custom GE 4×180k), which contained oligonucleotides ∼60 nucleotides (nt) in length comprised of: sequences within 100 bp of the center of HNF4α ChIP-seq peaks from proliferative Caco-2 cells (Verzi et al., 2010) were taken in 30-nt windows moving 5 nt at each step; 17,250 permutations of canonical HNF4α DR1 motifs (5’- AGGTCAAAGGTCA -3’); 500 permutations of DR2 motifs with variable spacer (5’- AGGTCNNNNGGTCA -3’); 900 random control 13-mer DNA sequences. A total of ∼45,000 test sequences were spotted in quadruplicate on the slide as single-stranded DNA. The DNA was made double-stranded and COS-7 or mouse liver NEs were applied. HNF4α binding was imaged with 2-µm resolution using Agilent G2565CA Microarray Scanner at the UCLA DNA Microarray Core. Extraction and normalization of the data were as described previously (Bolotin et al., 2010). PWMs were generated using seqLogo.

### Rapid Immunoprecipitation and Mass Spectrometry of Endogenous Proteins (RIME)

RIME was performed as previously described (Mohammed et al., 2016) with slight modifications. Livers from the same mice used for the RNA-seq and CHIP-seq -- WT and α7HMZ males n=3, 16-18 weeks of age sacrificed at 10:30 (ZT 3.5) or 20:30 (ZT13.5) -- were crosslinked and IP’d with the P1/P2 antibody. Multidimensional protein identification technology (MudPIT) analysis was performed by the UCR IIGB Proteomics Core. Raw MS1 and MS2 spectra were processed with Proteome Discoverer 2.1 (Thermo Scientific) and submitted to Mascot search engine to match against NCBI non-redundant mouse protein database. Only proteins with 1% FDR cut-off (q ≤ 0.01) were considered for subsequent analysis. Area under the curve, as reported by Proteome Discoverer, was averaged together for WT and α7HMZ samples (n=3) at each time point. IgG samples (n=3) from both WT and α7HMZ were averaged together to create a background sample. Areas were converted to log2 scale and the fold-change above IgG background was calculated for the WT and α7HMZ samples. Proteins with less than 8-fold change above background were omitted. Similarly, a 8-fold difference between WT and α7HMZ samples was used to identify unique protein interactions.

### Primary metabolite, complex lipids and oxylipin analysis

All metabolomic analysis was performed at the West Coast Metabolomics Center at the University of California Davis as described previously (Deol et al., 2017) using liver tissue rinsed in cold PBS, snap frozen and stored in liquid nitrogen. Data (pmol/gm tissue or peak height) are presented as mean +/- standard error of mean (SEM). Student’s T-test was used to determine statistical significance (p < 0.05) using GraphPad Prism v6.

Primary metabolite and complex lipid analysis was on WT (C57BL/6N) and α7HMZ (backcrossed into C67BL/6N) male mice harvested mid-morning and fed the standard chow (n=8, aged 38 weeks). Fold-enrichment was performed using MetaboAnalyst (Xia and Wishart, 2002). One outlier from each group was removed before plotting and statistical analysis. Analysis of non-esterified oxylipins was performed on a mixed 129/Sv plus C57BL/6 background WT and α7HMZ males (n=3 per group, aged 12-13 weeks). Tissue homogenates (100 mg) were extracted by solid phase extraction and analyzed by ultrahigh performance liquid chromatography tandem mass spectrometry (UPLC-MS/MS) (Agilent 1200SL-AB Sciex 4000 QTrap) as previously described (Matyash et al., 2008; Yang et al., 2009). Analyst software v.1.4.2 was used to quantify peaks according to corresponding standard curves with their corresponding internal standards.

## QUANTIFICATION AND STATISTICAL ANALYSIS

Differential gene expression (DEG) was measured using raw read counts with DESeq2: statistical significance was defined as adjusted p-value (padj) ≤ 0.01, unless otherwise noted. Legends denote thresholds using log2 fold change (log2FC) cutoffs. R library “gage” was utilized to identify differentially enriched KEGG pathways in Figure 1. Heatmaps were generated with pheatmap package in R; data were row-normalized before plotting, except for NR heatmap in Figure S2. Transcription Factor (TF) rankings for Cleveland plots were ordered at the 13:30 (peak HNF4α expression) then manually curated with the aid of PANTHER (Mi et al. 2017). Venn diagrams were generated with VennDiagram package in R. Unique and common RIME results were submitted to DAVID for ontology analysis. Statistical significance for primary metabolite and complex lipid data defined as p ≤ 0.05 by Mann-Whitney U-test or Benjamini-Hochberg padj <0.05, as indicated. All barplots represent mean ± SEM; significant differences are noted between genotypes at a given time point, unless indicated otherwise. For FPKM plots, padj values are from DESeq2; in other plots, p-values are from two-way Student’s T-test or One/Two-way ANOVA, as indicated. External expression datasets and analysis are described in Supplemental Methods.

## DATA AND SOFTWARE AVAILABILITY

The raw and processed RNA-seq data have been deposited in GEO under GSE117972.

The raw and processed ChIP-seq data have been deposited in GEO under (in progress).

The processed PBM data have been deposited in the Nuclear Receptor DNA Binding Project (http://nrdbs.ucr.edu).

The raw metabolomics data (primary metabolites and complex lipids) have been deposited in Metabolomics Workbench (www.metabolomicsworkbench.com) under Project #PR000461.

## Acknowledgements

We thank D Weaver for *Clock* KO mice, MC Weiss and N Briancon for HNF4α exon swap mice and J Vizcaya for assistance with newborn livers. The work was supported by NIH R01DK094707, DK053895 and USDA National Institute of Food and Agriculture (Hatch project CA-R-NEU-5680) to FMS; NIEHS T32 Training Grant (5T32ES018827) and Crohn’s and Colitis Foundation of America Career Development Award (#454808) to PD; WCMC Pilot Project from NIH U24 DK097154 to FMS, in collaboration with OF; R01ES002710 and Superfund Research Program P42 EX004699 to BDH; Start-up funds from UT Health to KEM; NIH S10 OD010669 for the Orbitrap Fusion mass spectrometer.

## Author Contributions

Conceptualization, J.R.D. and F.M.S.; Methodology, J.R.D., P.D., N.T., S.H.R., L.M.V., J.F., J.Y.; Software, J.R.D.; Validation, B.F., K.E-M.; Formal Analysis, J.R.D., P.D., J.F.; Investigation, J.R.D., P.D., N.T., S.H.R., L.M.V., J.R.E., S.P., J.F., J.Y., B.F.; Data Curation, J.R.D., P.D., J.F., F.M.S., Writing – Original Draft, J.R.D.; Writing – Review & Editing, J.R.D., P.D., N.T., L.M.V., K.E-M., F.M.S.; Visualization, J.R.D., P.D., S.H.R., B.F., K.E-M., F.M.S.; Supervision, B.D.H., O.F., K.E-M., F.M.S.; Project Administration, F.M.S.; Funding, B.D.H., O.F., K.E-M, F.M.S.

