## Supplemental Figures and Legends for "HNF4α isoforms regulate the circadian balance between carbohydrate and lipid metabolism in the liver"

**Deans et al.**

Deans_FigS1

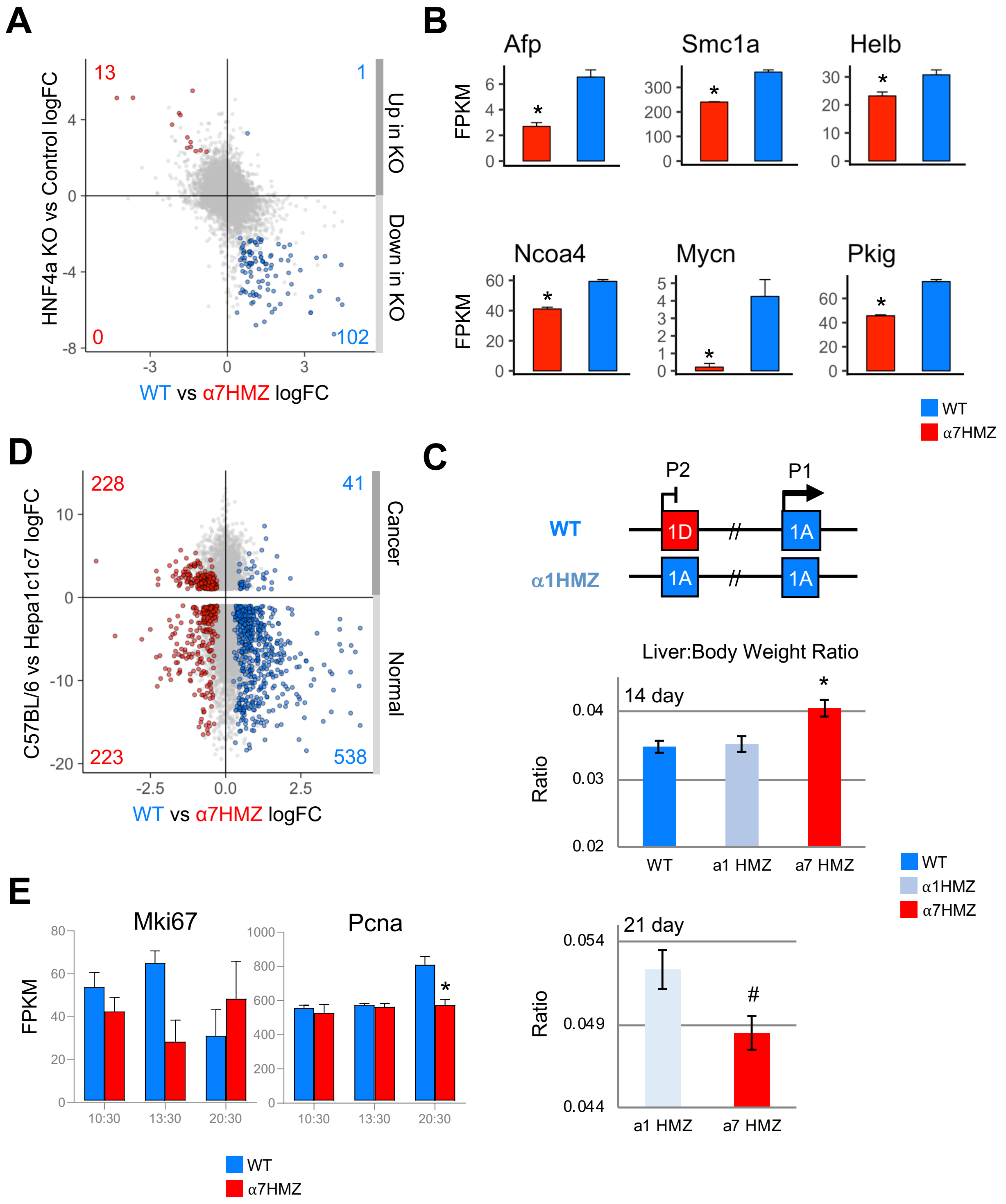
A
HNF4a KO vs Control logFC
WT vs α7HMZ logFC
Up in KO
Down in KO
13
1
0
102
B
Afp
Smc1a
Helb
Ncoa4
Mycn
Pkig
FPKM
WT
α7HMZ
C
P2
P1
WT
α1HMZ
1D
1A
Liver:Body Weight Ratio
14 day
21 day
Ratio
WT
a1 HMZ
a7 HMZ
α1HMZ
#
D
C57BL/6 vs Hepa1c1c7 logFC
WT vs α7HMZ logFC
Cancer
Normal
228
41
223
538
E
Mki67
Pcna
FPKM
10:30
13:30
20:30

**Figure S1 for Figure 1: HNF4α isoforms preferentially regulate genes in fetal liver and liver cancer.**

(*A*) Scatterplot of RNA-seq log2 fold-change (log2FC) values between WT and α7HMZ livers, plotted versus mouse liver HNF4α knockout (KO) microarray data. Colored data points with padj $\leq$ 0.01 in both datasets: blue dots, up in WT vs. α7HMZ; red dots, up in α7HMZ vs. WT. Numbers indicate highlighted genes in each quadrant. (*B*) FPKM barplots from RNA-seq of genes downregulated in α7HMZ, but more highly expressed in E14.5 fetal livers compared to adult. * padj $\leq$ 0.01. (*C*) Liver-to-body weight ratio of 14- and 21-day old mice (n=11 to 24). * p $\leq$ 0.01 One-way ANOVA 14-day; # p $\leq$ 0.05 Student's T-test 21-day. (*D*) as in (*A*) except WT and α7HMZ RNA-seq plotted vs. RNA-seq from murine hepatoma cell line (Hepa1c1c7) compared to non tumorigenic C57BL/6 control. (*E*) FPKM barplots of proliferation genes from RNA-seq. * padj $\leq$ 0.01. See Table S2AC for highlighted genes in (*A*) and (*D*).

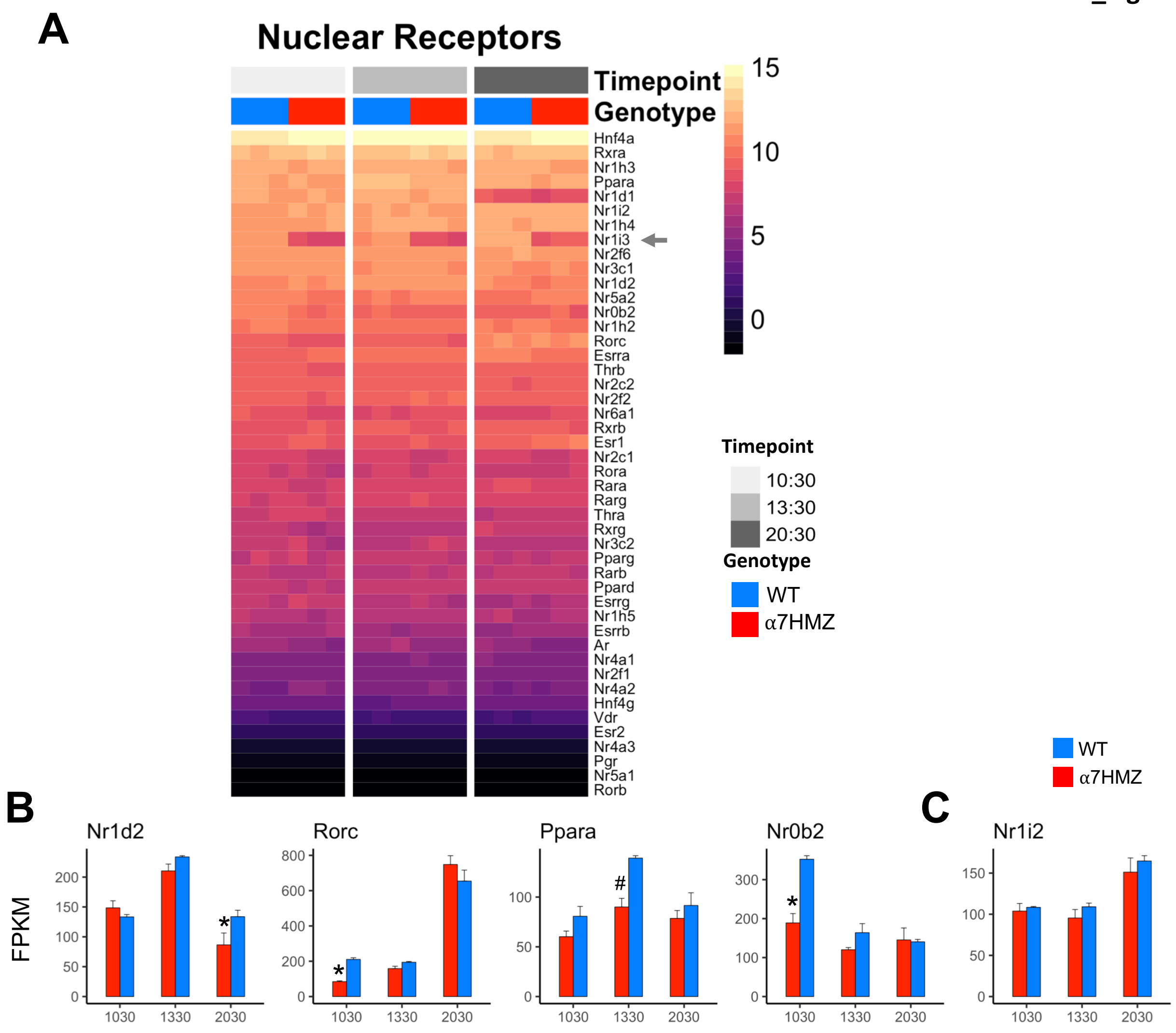


**Figure S2 for Figure 2: Dysregulation of nuclear receptors (NR) by HNF4α isoforms in the mouse liver.**

(*A*) Heatmap of regularized log-transformed (rlog) read counts for all NR in WT and α7HMZ males. Arrow, NR discussed in text. FPKM barplots of NR involved in the regulation of the circadian clock (*B*) and sex-specific expression of Cytochrome P450 genes (*C*) in WT and ***α***7HMZ males. * padj $\leq$ 0.01; # padj $\leq$ 0.05

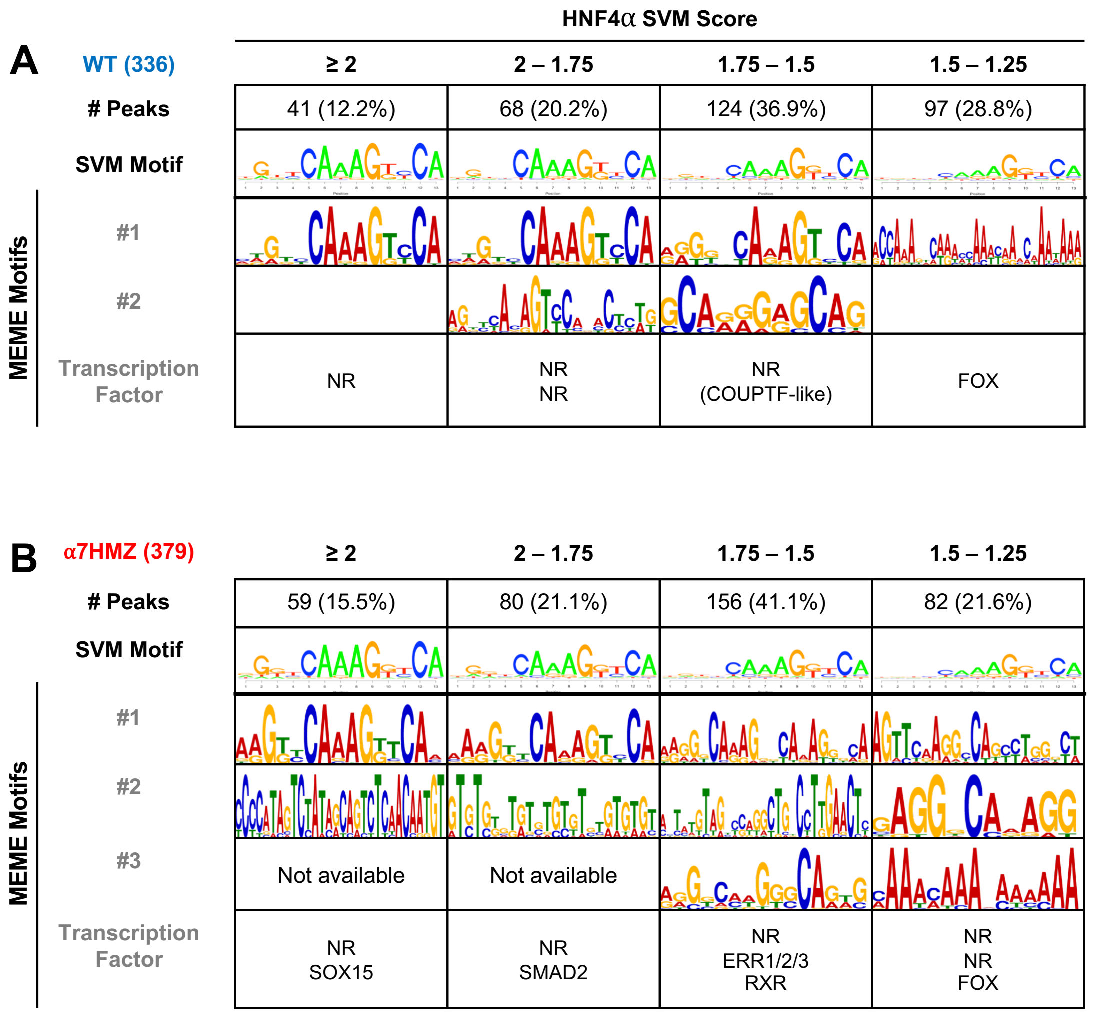


**Figure S3 for Figure 4. Motif analysis of HNF4α ChIP-seq peaks categorized by HNF4α motifs.**

Categorization of WT- and α7HMZ-unique ChIP peaks into four groups based on highest SVM HNF4α motif score, as noted. Number of unique peaks for each genotype are given in parentheses. HNF4α SVM-derived binding motifs are shown. Top motifs derived from *de novo* MEME-ChIP analysis are shown with TF corresponding to the motifs listed below. NR, HNF4α-like DR1 motif.

Deans_FigS4

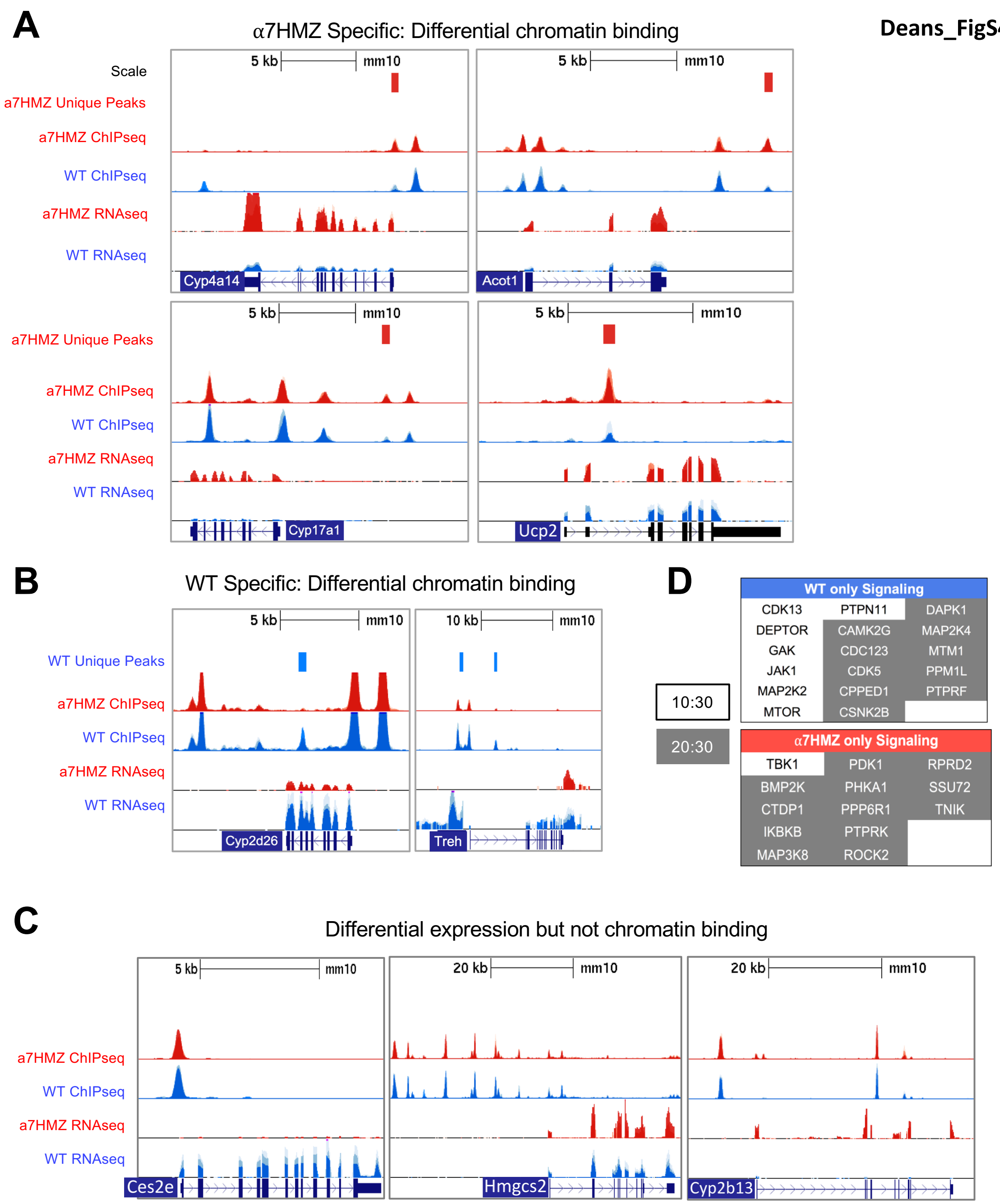
A
α7HMZ Specific: Differential chromatin binding
Scale
5 kb
mm10
a7HMZ Unique Peaks
a7HMZ ChIPseq
WT ChIPseq
a7HMZ RNAseq
WT RNAseq
Cyp4a14
Acot1
Cyp17a1
Ucp2
B
WT Specific: Differential chromatin binding
10 kb
WT Unique Peaks
Cyp2d26
Treh
C
Differential expression but not chromatin binding
20 kb
Ces2e
Hmgcs2
Cyp2b13
D
10:30
20:30
WT only Signaling
CDK13
PTPN11
DAPK1
DEPTOR
CAMK2G
MAP2K4
GAK
CDC123
MTM1
JAK1
CDK5
PPM1L
MAP2K2
CPPED1
PTPRF
MTOR
CSNK2B
α7HMZ only Signaling
TBK1
PDK1
RPRD2
BMP2K
PHKA1
SSU72
CTDP1
PPP6R1
TNIK
IKBKB
PTPRK
MAP3K8
ROCK2

**Figure S4 for Figure 5: Examples of dysregulated genes with unique ChIP-seq peaks.**

(*A*) UCSC Genome Browser view of four α7HMZ genes with a unique ChIP-signal within ~10 kb of TSS. ChIP-seq and uniquely bound regions are in the top two tracks; RNA-seq from 10:30 AM is in the bottom two tracks. (*B*) as in (*A*) View of two WT genes with unique ChIP signal within ~10 kb of TSS. (*C*) RIME results showing proteins involved in signaling pathways uniquely bound to HNF4α in WT or α7HMZ livers at 10:30 AM and 20:30.

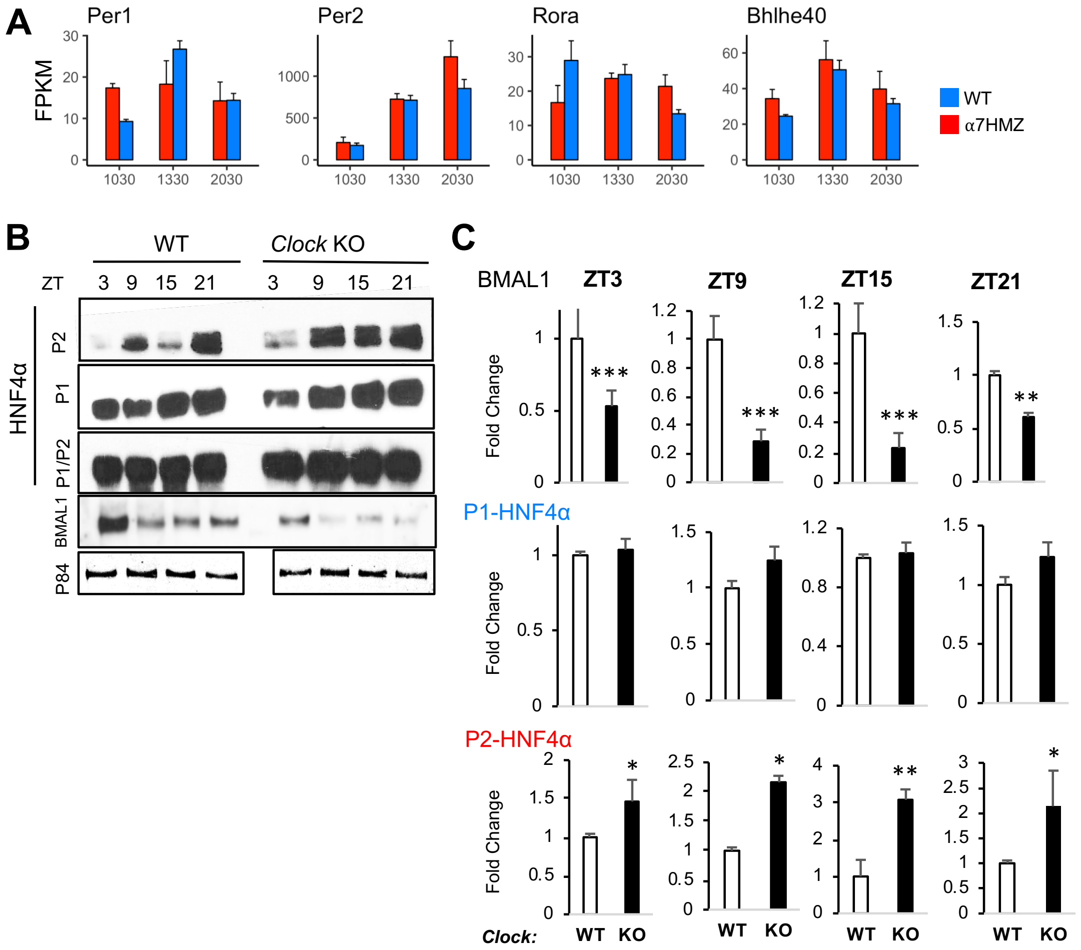


**Figure S5 for Figure 6: HNF4α isoforms impact the circadian response and are regulated by the clock.**

(*A*) Barplots of FPKM values for core circadian TFs not significantly dysregulated (padj $\geq 0.05$) in WT and α7HMZ livers. (*B*) Representative immunoblots showing levels of P1- and P2-HNF4α protein in WT and *Clock* KO mouse liver whole cell extracts at the indicated time points, using antibodies that recognize either a single isoform (P1 or P2) or both isoforms (P1/P2). Also shown are BMAL1 (*Arntl*) and P84. (*C*) Quantification of BMAL1, P1-HNF4α and P2-HNF4α protein from immunoblots of individual livers (n=3-4) from WT and *Clock* KO mice throughout the circadian cycle. * $p \leq 0.01$; ** $p \leq 0.001$; *** $p \leq 0.001$ by two-tailed Student's T-test.

Deans_FigS6

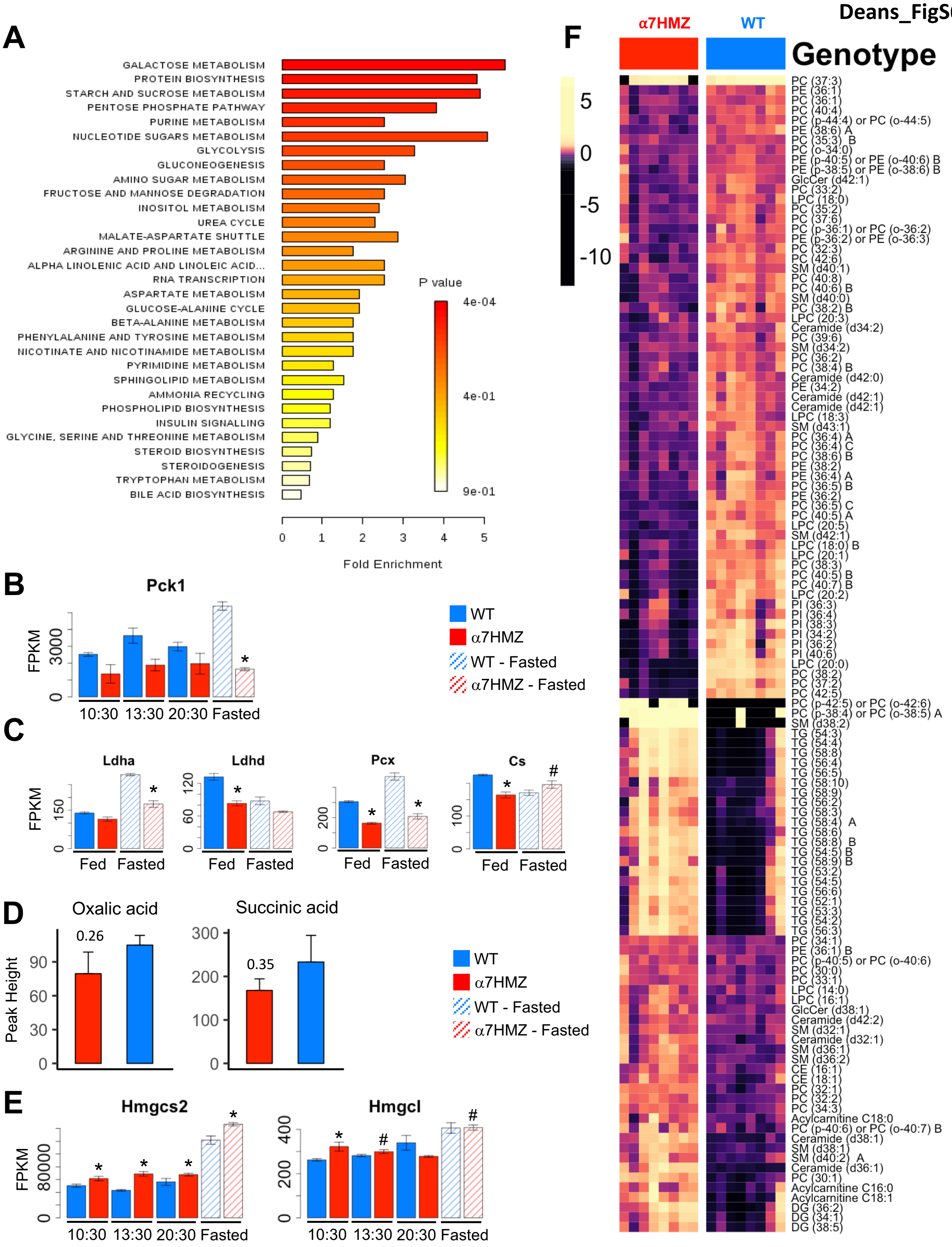
A
GALACTOSE METABOLISM
PROTEIN BIOSYNTHESIS
STARCH AND SUCROSE METABOLISM
PENTOSE PHOSPHATE PATHWAY
PURINE METABOLISM
NUCLEOTIDE SUGARS METABOLISM
GLYCOLYSIS
GLUCONEOGENESIS
AMINO SUGAR METABOLISM
FRUCTOSE AND MANNOSE DEGRADATION
INOSITOL METABOLISM
UREA CYCLE
MALATE-ASPARTATE SHUTTLE
ARGININE AND PROLINE METABOLISM
ALPHA LINOLENIC ACID AND LINOLEIC ACID...
RNA TRANSCRIPTION
ASPARTATE METABOLISM
GLUCOSE-ALANINE CYCLE
BETA-ALANINE METABOLISM
PHENYLALANINE AND TYROSINE METABOLISM
NICOTINATE AND NICOTINAMIDE METABOLISM
PYRIMIDINE METABOLISM
SPHINGOLIPID METABOLISM
AMMONIA RECYCLING
PHOSPHOLIPID BIOSYNTHESIS
INSULIN SIGNALLING
GLYCINE, SERINE AND THREONINE METABOLISM
STEROID BIOSYNTHESIS
STEROIDOGENESIS
TRYPTOPHAN METABOLISM
BILE ACID BIOSYNTHESIS
P value
4e-04
4e-01
9e-01
0 1 2 3 4 5
Fold Enrichment
B
Pck1
FPKM
0 3000
10:30 13:30 20:30 Fasted
WT
α7HMZ
WT - Fasted
α7HMZ - Fasted
C
Ldha
Ldhd
Pcx
Cs
FPKM
Fed Fasted
D
Oxalic acid
Succinic acid
Peak Height
0.26
0.35
0 30 60 90
0 100 200 300
E
Hmgcs2
Hmgcl
FPKM
0 80000
0 200 400
10:30 13:30 20:30 Fasted
F
α7HMZ
WT
Genotype
5
0
-5
-10
PC (37:3)
PE (36:1)
PC (36:1)
PC (40:4)
PC (p-44:4) or PC (o-44:5)
PE (38:6) A
PC (35:3) B
PC (o-34:0)
PE (p-40:5) or PE (o-40:6) B
PE (p-38:5) or PE (o-38:6) B
GlcCer (d42:1)
PC (33:2)
LPC (18:0)
PC (35:2)
PC (37:6)
PC (p-36:1) or PC (o-36:2)
PE (p-36:2) or PE (o-36:3)
PC (32:3)
PC (42:6)
SM (d40:1)
PC (40:8)
PC (40:6) B
SM (d40:0)
PC (38:2) B
LPC (20:3)
Ceramide (d34:2)
PC (39:6)
SM (d34:2)
PC (36:2)
PC (38:4) B
Ceramide (d42:0)
PE (34:2)
Ceramide (d42:1)
Ceramide (d42:1)
LPC (18:3)
SM (d43:1)
PC (36:4) A
PC (36:4) C
PC (38:6) B
PE (38:2)
PE (36:4) A
PC (36:5) B
PE (36:2)
PC (36:5) C
PC (40:5) A
LPC (20:5)
SM (d42:1)
LPC (18:0) B
LPC (20:1)
PC (38:3)
PC (40:5) B
PC (40:7) B
LPC (20:2)
PI (36:3)
PI (36:4)
PI (38:3)
PI (34:2)
PI (36:2)
PI (40:6)
LPC (20:0)
PC (38:2)
PC (37:2)
PC (42:5)
PC (p-42:5) or PC (o-42:6)
PC (p-38:4) or PC (o-38:5) A
SM (d38:2)
TG (54:3)
TG (54:4)
TG (58:8)
TG (56:4)
TG (56:5)
TG (58:10)
TG (58:9)
TG (56:2)
TG (58:3)
TG (58:4) A
TG (58:6)
TG (58:8) B
TG (54:5) B
TG (58:9) B
TG (53:2)
TG (54:5)
TG (56:6)
TG (52:1)
TG (53:3)
TG (54:2)
TG (56:3)
PC (34:1)
PE (36:1) B
PC (p-40:5) or PC (o-40:6)
PC (30:0)
PC (33:1)
LPC (14:0)
LPC (16:1)
GlcCer (d38:1)
Ceramide (d42:2)
SM (d32:1)
Ceramide (d32:1)
SM (d36:1)
SM (d36:2)
CE (16:1)
CE (18:1)
PC (32:1)
PC (32:2)
PC (34:3)
Acylcarnitine C18:0
PC (p-40:6) or PC (o-40:7) B
Ceramide (d38:1)
SM (d38:1)
SM (d40:2) A
Ceramide (d36:1)
PC (30:1)
Acylcarnitine C16:0
Acylcarnitine C18:1
DG (36:2)
DG (34:1)
DG (38:5)

**Figure S6 for Figure 7: Metabolic effects of P2-HNF4α in the adult liver.**

(*A*) Pathway enrichment for known primary metabolites dysregulated between WT and α7HMZ livers, with a Mann-Whitney U-test $p \leq 0.05$. (*B,C,E*) FPKM values for the indicated genotypes and time points (Fed in *C* is 10:30 AM) * padj $\leq 0.01$; # padj $\leq 0.05$. (*D*) Bar plots for Krebs cycle metabolites trending towards repression in α7HMZ, but not statistically significant. (*F*) Heatmap of row normalized levels for known complex lipids in WT and α7HMZ livers (Benjamini-Hochberg padj $\leq 0.05$).

Deans_FigS7

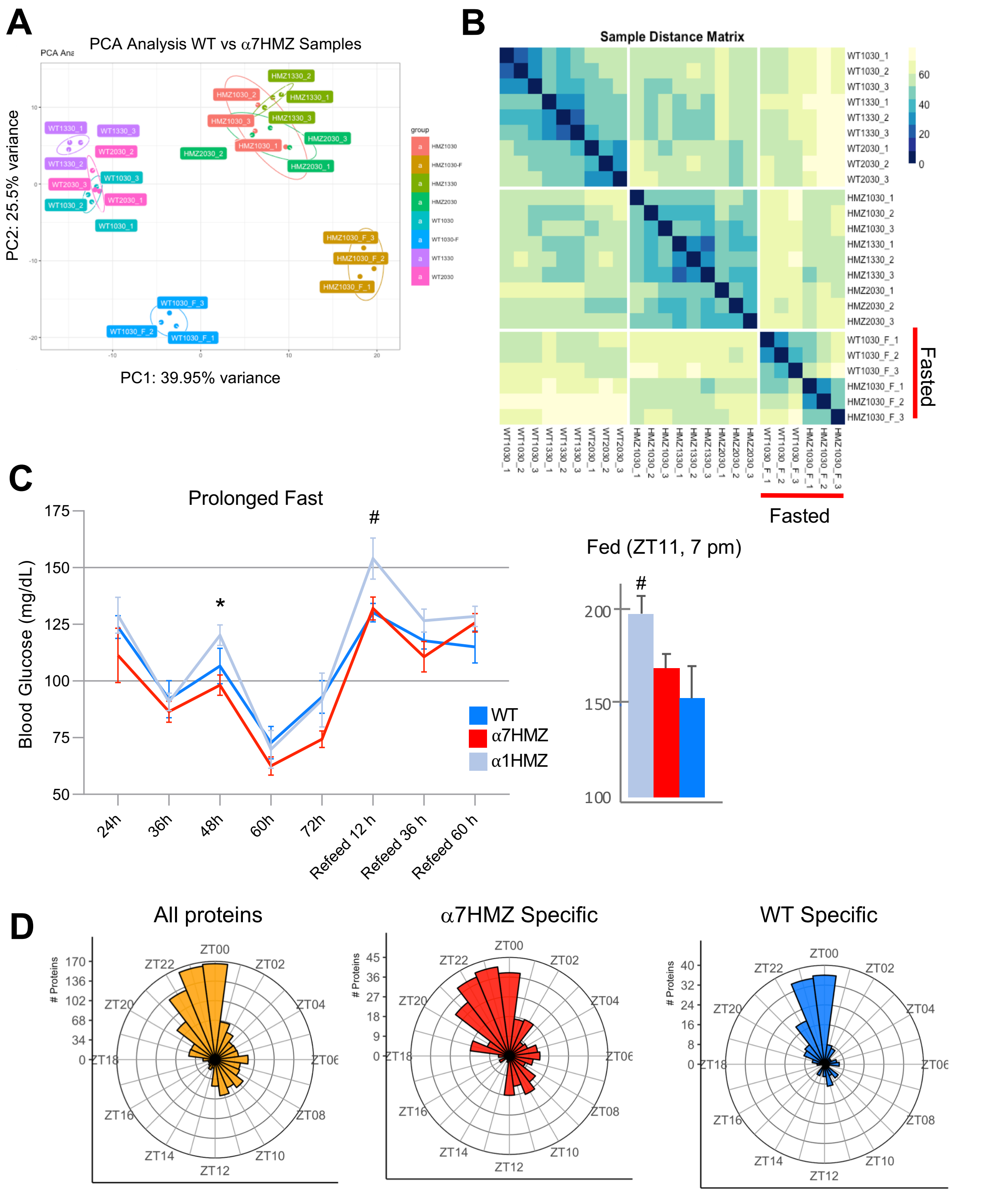
A
PCA Analysis WT vs α7HMZ Samples
PC2: 25.5% variance
PC1: 39.95% variance
B
Sample Distance Matrix
Fasted
C
Prolonged Fast
Blood Glucose (mg/dL)
WT
α7HMZ
α1HMZ
24h
36h
48h
60h
72h
Refeed 12 h
Refeed 36 h
Refeed 60 h
Fed (ZT11, 7 pm)
D
All proteins
α7HMZ Specific
WT Specific
### Proteins

**Figure S7 for Figure 7: PCA and Sample Distance Matrix for all RNA-seq samples.**

(*A*) PCA analysis of all RNA-seq samples using distance matrix for entire transcriptome. Biological replicates denoted by underscores and digit, fasting samples denoted by "_F". (*B*) Sample Distance Matrix for each RNA-seq replicate, including fasting, calculated for entire transcriptome. Dark blue indicates smaller distance which implies high degree of similarity. (C) *Left,* circulating blood glucose levels of male mice (~6 mo. old) starting at 24 hr after food was removed (ZT11). Mice were re-fed after 72 hr of fasting. WT and α1HMZ n=5; α7HMZ n= 6 for 24, 36 and 48 hr, n=4 for 60 hr, n=3 for remaining time points. #, $p< 0.05$ α1HMZ vs. WT; * $p<0.01$α1HMZ vs. α7HMZ. *Right*, blood glucose levels from fed males (3 to 6 mo. old) at ZT11. WT n=9, α1HMZ n =10, α7HMZ n= 7. #, $p< 0.05$ α1HMZ versus WT. $p=0.055$ for α1HMZ vs. α7HMZ. (*D*) Rose plots of All Proteins from Robles et al., 2014 (*left*) filtered for DEGs at any time point (10:30, 13:30, 20:30, padj ≤ 0.05) specific for α7HMZ (*middle*) or WT (*right*).
