## Supplemental Materials for "HNF4α isoforms regulate the circadian balance between carbohydrate and lipid metabolism in the liver"

### SUPPLEMENTAL METHODS

#### EXPERIMENTAL MODEL AND SUBJECT DETAILS

#### Cell Lines

COS-7 cells (ATCC CRL 1651) were maintained at 37°C and 5% $CO_2$ in Dulbecco's modified Eagle's medium (DMEM) supplemented with 10% bovine calf serum (BCS).

#### Animal Models

WT and α7HMZ mice were maintained in isolator cages under 12-h light/dark cycles at ~21$^0$C on bedding (Andersons bed OCOB Lab ⅛ 1.25CF) from Newco (Rancho Cucamonga, CA) and fed a standard lab chow (LabDiet, #5001, St. Louis, MO). They were bred and maintained in a specific pathogen free (SPF) vivarium. All experiments were performed in an SPF/barrier vivarium, except for the fasting glucose/ketone body experiments for which the mice were housed in a conventional vivarium with mice maintained in individually ventilated cages. Adult males were used for all experiments except the newborn liver analysis, for which gender was not identified, and the female mice in Figure 7H. Wood chip bedding was used for all fasting experiments.

The transgenic mice on a mixed 129/Sv plus C57BL/6 background carrying exon 1A or exon 1D in both the P1 and P2 promoter (α1HMZ and α7HMZ, respectively) have been described previously (Briançon and Weiss, 2006). Both lines were maintained as heterozygotes; wildtype (WT) and homozygous (α7HMZ) were mated for a single generation to generate mice for the experiments. The WT and α7HMZ mice in the mixed background were used for all RNA-seq, ChIP-seq, RIME and oxylipin analysis experiments. Mice backcrossed 10+ generations into C57BL/6N were used for all metabolomic and ketone body/glucose experiments. Mice of the same genotype were housed 3-5 per cage and randomly selected to treatment groups at the beginning of the experiment. Mice were euthanized by $CO_2$ asphyxiation and tissues harvested at the designated experiment time points. All mice used were adult males, aged 16 to 20 weeks, unless otherwise noted. Time points were 10:30 (ZT 3.5), 13:30 (ZT 6.5) and 20:30 (ZT 13.5) (lights on at 7:00 and off at 19:00).

The α7HMZ mice used for metabolomic analysis were backcrossed to C57BL/6N for 10+ generations and used with C57BL/6N WT controls (n=8, 35 weeks of age). They were fed the standard lab chow and sacrificed mid morning. For the newborn liver analysis, α7HMZ and α1HMZ (both backcrossed 10+ generations into C57BL/6N) were compared to C57BL/6N (WT) controls 14 or 21 d after birth. No distinction between males and females were made.

*Clock*-deficient (Clock KO) mice were provided by Dr. David Weaver and have been described previously (Debruyne et al., 2006). The *Clock* KO colony was maintained and propagated by mating heterozygous Clock+/- male and female mice in order to generate *Clock* KO and WT littermate pups. Mice were fed *ad libitum* and kept in 12-h light/dark cycle in a semi-barrier, SPF vivarium in individually ventilated cages. WT and *Clock* KO mice were sacrificed at the indicated zeitgeber times (ZT) and livers were flash frozen in liquid nitrogen for future analysis by immunoblot and qPCR.

Tail bleeds from live mice were used for glucose measurements with a One Touch Ultra glucose meter and for ketone body measurements with a Precision Ketone meter, except for the 60-hr fast in Figure 7G which used blood from a post-mortem cardiac puncture analyzed by a β-Hydroxybutyrate (Ketone Body) Colorimetric Assay Kit from Cayman (Item No. 700190).

Care and treatment of the animals were in strict accordance with guidelines from the Institutional Animal Care and Use Committee at the University of California, Riverside, or the McGovern Medical School, UTHealth.

**METHOD DETAILS**

*RNA Extraction and Reverse Transcriptase, Quantitative PCR*

Total RNA was isolated from cells and liver using TRIzol reagent (Thermo Scientific) according to the manufacturer's protocol. One µg of total RNA was used for cDNA synthesis using an iScript cDNA synthesis kit (BioRad, #1708891). Advanced Universal SYBR Green Supermix (BioRad, #4385618) was used for qPCR amplification using a Bio-Rad C1000. PCR protocol settings were as follows: 95°C for 30 sec, 95°C for 10 sec, 62°C for 30 sec and then 39 cycles at 65°C for 31 sec and 65°C for 5 sec. 18S RNA was used as control. The fold change in mRNA expression for each gene was calculated using $2^{-\Delta\Delta C}$.

*Immunoblot (IB) analysis*

Immunoblots (IBs) in Figure 1 were carried out as previously described (Jiang et al., 1995). Proteins from nuclear extracts (NE) and whole cell extracts (WCE) were separated by 10% SDS-PAGE and then transferred to PVDF (Immobilon). Even loading was verified by Coomassie stain of the blot. Primary antibodies (Ab) were mouse monoclonal anti-HNF4α P1/P2 (R&D Systems #PP-H1415-00) which recognizes the C-terminus of both P1- and P2-HNF4α isoforms, and mouse monoclonal anti-HNF4α P1 (R&D Systems # PP-K9218-00) which recognizes the N-terminus of P1-HNF4α. Both were used at 1:10,000 overnight. Secondary antibodies were horseradish peroxidase (HRP)-conjugated goat anti-mouse (GαM-HRP) Abs from Jackson ImmunoResearch Laboratories. Development was with Dura Kit (Fisher Scientific #34075). The procedure for NE from COS-7 and liver are described below. WCE of liver in Figure 1 were prepared using RIPA buffer (see RIME below).

For *Clock* KO and WT liver IBs, approximately 100 mg of liver tissue was homogenized in modified RIPA buffer: 50 mm Tris-HCl pH 8.0, 150 mM NaCl, 5 mM EDTA, 15 mM $MgCl_2$, 1% NP-40, 1 mM PMSF, 1mM NaF, 400 mM NAM (nicotinamde, sirtuin inhibitor) and 3.3 mM TSA (trischostatin A, HDAC inhibitor) and protease inhibitors (Roche, #11697498001) and using a MagNA Lyser (Roche). Samples were spun at high speed for 10 min to remove insoluble material (Thermoscientific Sorvall ST16R, 10,000 rpm with rotor F21-48/2 for 9520 RCF). The total protein levels of the lysates were determined using the Pierce BCA Protein Assay Kit (Pierce, #23225). Twenty micrograms of whole cell lysate was analyzed using 8% SDS-PAGE followed by transfer to nitrocellulose membranes before staining with primary and secondary antibodies. Primary antibodies included P1/P2-HNF4α (#PP-H1415-00 R&D Systems) at 1:1000, P1-HNF4α (#PP-K9218-00 R&D Systems) at 1:1000, P2-HNF4α (#PP-H6939-00 R&D Systems) at 1:1000, BMAL1 (#93806 Abcam) at 1:1000, and p84 (# GTX70220 GeneTex). Secondary antibodies included Anti-Mouse-HRP (#1706516 BioRad) at 1:15,000 and Anti-Rabbit-HRP (#1791019 BioRad), also at 1:15,000). Blots were visualized with Clarity Chemiluminescence Reagent (ECL) (Biorad, #170-5061).

*Preparation of nuclear extracts (NE) for Immunoblot (IB) and Protein Binding Microarrays (PBM) analysis*

Following buffers and inhibitors were used:

**TE:** 10 mM Tris-HCl, 1 mM EDTA pH 8.0

**2X HBS:** 274 mM of NaCl, 10 mM KCl, 1.4 mM Na2HPO4, 15 mM D-glucose, 42 mM HEPES (free acid), pH 7

**1X H Buffer**: 100 mM HEPES, pH 7.8, 250 mM KCl, 1.5 mM spermine, 5 mM spermidine, 10 mM EGTA, 10 mM EDTA

**Buffer A**: 1X H Buffer, 0.32 M Sucrose

**Low salt buffer**: 1X H buffer, 20% glycerol

**High salt buffer**: 1X H buffer, 20% glycerol, 1 M KCL (for COS-7 cells) or 0.5 M KCL (for liver).

**Inhibitors and DTT:** Protease inhibitors (Sigma, #8340), Phosphatase inhibitor cocktail I (Sigma, #P-2850), Phosphatase inhibitor cocktail II (Sigma, #P-5726) and 200 mM phenylmethylsulfonyl fluoride (PMSF) were added at the dilution 1:200 (for cell NE) and 1:100 (for liver NE) to each buffer solution before each use. Dithiothreitol (DTT) was added to 1 mM final.

Nuclear extracts (NE) were prepared from COS-7 cells transiently transfected via $CaPO_4$ with HNF4α expression vectors for human HNF4α2 (NM_00457) and HNF4α8 (NM_175914) as previously described (Jiang et al., 1995) with some modifications. Cells (3.5 x $10^6$) were plated in 150-mm plates and incubated in 15 ml of DMEM supplemented with 10% BCS at 37°C for 24 to 48 hours. Twenty-five µg of plasmid DNA, HNF4α2 (NM_00457) or HNF4α8 (NM_175914) in pcDNA3.1, was mixed with 450 µl TE and 500 µL 2X HBS buffer; 50 µL 2.5 M $CaCl_2$ was added to the mixture, and 25 min later the mixture was added to the cells. After approximately 10 h of incubation, the cells were washed 1x with PBS and then shocked with 3 mL 15% glycerol in 1X HBS buffer for 3 min 15 s, and then washed 2X with PBS, followed by DMEM plus 10% BCS. Then, 24 to 32 h later, the cells were harvested and NE were prepared. Cells were washed twice with cold PBS, then once with 1 ml of 0.25X Buffer H; 0.75 ml of 0.25X Buffer H was subsequently added to each 150-mm plate and incubated on ice for 5 to 20 min. Cells were scraped and resuspended in equal volume of 2X Buffer H plus 20% glycerol. After centrifugation (10 min at 2,500 rpm) the supernatant was discarded and the nuclear pellet resuspended in an equal volume of Low Salt Buffer; 0.72X volumes of High Salt Buffer was used to resuspend the nuclei followed by nutation at 4°C for 1h 10 min. The soluble NE was separated from the chromatin pellet by centrifugation (25 min at 12,000 rpm). Samples were snap-frozen and subsequently used for IBs and PBMs.

Liver NE from WT and α7HMZ mice were prepared as previously described (Yuan et al., 2009) by motorized homogenization of frozen or fresh liver in Buffer A plus 0.3% Triton X-100, protease and phosphatase inhibitors. The homogenate was filtered using 100-μm cell strainers (Fisher #08-771-19) before passing through the dounce homogenizer (Fisher #06-435B) in Buffer A plus 0.3% Triton X-100 followed by nuclei separation via centrifugation (10 min at 3,300 rpm) and multiple washes of nuclei (1X wash in Buffer A plus 0.3% Triton X-100, followed by 2X wash in Low Salt Buffer). Each wash was followed by centrifugation (10 min at 2,000 rpm). After washes, nuclear pellets were resuspended in 1X volume of Low Salt Buffer, 2X volume of High Salt Buffer was added, and extraction was performed as described above for cell NE. All incubations, separations and washes were at 4°C.

*Expression profiling (RNA-seq) and Analysis*

Next generation sequencing of RNA (RNA-seq) was carried out as previously described (Vuong et al., 2015). WT and α7HMZ male mice were sacrificed (n=3, aged 16-18 weeks) at each time point: 10:30, 13:30, 20:30 PM (ZT 3.5, ZT 6.5, and ZT 13.5, respectively). The three mice were harvested in succession within a 30-min time frame. Two ~25 mg pieces from each liver were immediately frozen in liquid nitrogen and stored at -4°C. The miRNeasy Mini Kit (Qiagen, #74104) was used to extract and purify total RNA; 4 μg of each sample was used to prepare a poly(A)+ RNA library using TruSeq RNA Sample Prep v2 Kit (Illumina, Cat# RS-122-2001). Libraries submitted for 75-bp single-end sequencing with Illumina NextSeq 500 at the UCR IIGB Genomics Core. A total of 24 libraries (3 fed time points, 1 fasted time point, 2 genotypes each, 3 replicates) were multiplexed and sequenced in two separate runs, each of which yielded ~600 M reads, averaging ~50 M reads per sample.

Reads were aligned to the mouse reference genome, mm10, with Illumina's iGenome genes.gtf file using TopHat v2.1.1 using default parameters with the exception of allowing only 1 unique alignment for a given read. Raw read counts were calculated at the gene level for each sample using HTSeq v0.6.1. Library normalization was performed with EDASeq; within-lane normalization on GC content was performed with the LOESS method and between-lane normalization was performed with non-linear full quantile method. Normalization factors from EDASeq were used for differential expression analysis with DESeq2. Normalized read counts, FPKM (fragments per kilobase per million), and rlog (regularized log transformation) results were generated for downstream analysis. Pairwise contrasts were generated for all relevant comparisons. Sample distance matrix were generated using rlog transformed values from DESeq2.

*Chromatin Immunoprecipitation Sequencing (ChIP-seq) and Analysis*

A freshly minced chunk from the large lobe of the liver (approximately 100-200 mg) of a 16 to 20-week old male mouse (WT and α7HMZ, 10:30) was fixed in 1% formaldehyde (in ChIP Buffer: 1X PBS plus 1 mM PMSF, 1 mM DTT, 2 μg/mL leupeptin, 2 μg/mL aprotinin) for 15 min at room temperature (RT). The crosslink reaction was stopped with 0.125 M glycine for 5 min at RT and centrifuged (10 min at 2,000 rpm) at 4°C. All subsequent steps were performed at 4°C. Fixed tissue was further processed using motorized and glass dounce homogenizers (Fisher #06-435B) in cold ChIP Buffer. The homogenate was filtered using 100-μm cell strainers (Fisher #08-771-19) before passing through the dounce homogenizer. Isolated liver cells were processed as previously described (Vuong et al., 2015). Briefly, cells were swelled in 1.0 mL cold Hypotonic Buffer (10 mM HEPES-KOH pH 7.9, 10 mM KCl, 1.5 mM $MgCl_2$) plus 1 mM PMSF and 1 mM DTT) for 10 min. The nuclei were collected by centrifugation and resuspended in 1.0 mL cold Nuclei Lysis Buffer (1% SDS, 50 mM Tris-HCl pH 8.0, 10 mM EDTA) plus 1 mM PMSF, 1 mM DTT, 2 µg/mL leupeptin and 2 µg/mL aprotinin. The samples were sonicated using a Sonic Dismembrator Model 500 (Fisher Scientific) to obtain DNA fragments of about 200-500 bps, diluted 1:1 with Immunoprecipitation (IP) dilution buffer (0.01% SDS, 20 mM Tris-HCl pH 8.0, 1.1% Triton X-100, 167 mM NaCl, 1.2 mM EDTA), and pre-cleared with 20 µL of packed Protein G Agarose (Pierce) beads (1:1 slurry in IP dilution buffer) that were pre-blocked with 100 µg/µL BSA for 30 min. The IP was performed with 4.2 µg of affinity-purified anti-HNF4α (α-445) (Sladek et al., 1990) or rabbit IgG control (Santa Cruz, cat#sc-2027). Thirty to forty microliters of packed protein G beads (1:1 slurry) were added to the IP sample and incubated overnight. All IPs were washed with three sequential buffers for 5 min each at RT: TSE I (0.1% SDS, 1% Triton X-100, 2 mM EDTA, 20 mM Tris-HCl pH 8.0, 150 mM NaCl), TSE II (as TSE I but with 500 mM NaCl) and TSE III (0.25 mM LiCl, 1% NP-40, 1% Deoxycholate, 1 mM EDTA, 10 mM Tris-HCl pH 8.0). At the final wash, the IP sample was washed twice with 1X TE for 5 min at RT. The precipitated material was then eluted with IP elution buffer (1% SDS and 0.1 M $NaHCO_3$) twice. For the first elution, the sample was eluted with 150 µL buffer, incubated at RT for 20 min, centrifuged at maximum (max) speed for 1 min and the supernatant was transferred to a new tube. The second elution was the same as the first but with an additional 1 min boiling in a 100°C heat block preceding the 20-min incubation. The material from the first and second elutions were combined, incubated at 65°C for 4 to 5 h to reverse the crosslinks. DNA precipitation was performed with 1 mL of 100% ethanol overnight at -20°C. Protein and RNA digestions were performed for 1 h at 55°C with 11 μL of 10X Proteinase K buffer (100 mM Tris-HCl pH 8.0, 50 mM EDTA, 500 mM NaCl) and 1 μL of 19 mg/mL Proteinase K solution (in 10 mM Tris-HCl pH 8.0, 1 mM $CaCl_2$ plus 30% glycerol) and 25 min at RT with 1 μL of 10 μg/uL RNaseA in dd$H_2O$, respectively. DNA material was purified with GeneJET PCR Purification Kit (Thermo Scientific, #K0701). Qubit fluorometer at the University of California Riverside (UCR) Institute for Integrative Gene Biology (IIGB) Genomics Core was used to measure DNA concentration; 2-22 μg of ChIP'd material was used to generate a library using the BIOO Scientific ChIPseq DNA Library Kit. Libraries were submitted for 50-bp single end sequencing by Illumina HiSEQ 2500 at the IIGB core facility.

Reads were aligned to the mouse reference genome, version mm10, with Bowtie2 using the default parameters. Peaks were called with MACS2 using default parameters for individual samples, as well as a pooled peak dataset using the SPMR (signal per million reads) parameter. Aligned reads and MACS2 peak-sets were analyzed with DiffBind to identify common and uniquely bound regions of the genome. Livers from three mice were used for each genotype. After PCA analysis, one α7HMZ replicate was identified visually as an outlier from the other two replicates so a single WT and α7HMZ replicate were omitted from downstream analysis, leaving two replicates per genotype. DiffBind analysis was performed with default parameters using DESeq2 for analysis and library size calculated as total aligned reads. ChIP peaks were excluded from analysis if MACS2 results showed -log10(p-value) ≤ 10 to produce a filter removing peaks below six-fold enrichment over background, and unique peak IDs were manually annotated to each resulting peak. Manually curated peak lists were generated by filtering all results on peaks with "concentration" ≥ 5.5. Concentration defined by DiffBind as the "mean (log) reads across all samples" in contrast.

*Support Vector Machine (SVM) Predictions and Motif Generation*

The kernel-based SVM was trained as previously described using results from independent HNF4α PBM experiments (Bolotin et al., 2010). All possible 13-mers in both orientations from each uniquely bound ChIP peak were submitted to the HNF4α PBM SVM (http://nrmotif.ucr.edu/fuzzhtmlform.html) for score predictions using Kernlab package in R. Each ChIP peak was then annotated with the score and sequence of the single highest predicted motif from all possible k-mers and categorized into four different bins. All sequences within each bin were submitted to seqLogo (Bembom 2017) to generate a position weight matrices (PWM) representing the strongest HNF4α binding motif within the category. For each SVM-score category, sequences from a 200-bp window around the peak center for each ChIP peak were submitted to MEME-ChIP (Machanick and Bailey, 2011) for *de novo* motif analysis with default parameters, with the exception of number of motifs to identify (6), max word size (24), and the transcription factor binding site (TFBS) database utilized was HOCOMOCO v10.

*Protein Binding Microarrays (PBM)*

Protein binding microarrays (PBMs) were carried out as previously described (Bolotin et al., 2010). A custom-designed array was ordered from Agilent (SurePrint G3 Custom GE 4x180k), which contained oligonucleotides ~60 nucleotides (nt) in length, corresponding to the following sequences: sequences within 100 bp of the center of published HNF4α ChIP-seq peaks from proliferative Caco-2 cells (Verzi et al., 2010) were taken in 30-nt windows moving 5 nt at each step; 17,250 permutations of canonical HNF4α DR1 motifs (5’- AGGTCAAAGGTCA -3’); 500 permutations of DR2 motifs with variable spacer (5’- AGGTCNNNNGGTCA -3’); 900 random control 13-mer DNA sequences. A total of ~45,000 test sequences were spotted in quadruplicate on the slide as single-stranded DNA. The DNA was extended and made double-stranded on the slide using a primer to a common linker sequence (5'-TCGACCGATACTCTAATCTCCCTAGGC-3'), dNTPs (GE Healthcare) and Thermo Sequenase (Affymetrix, Cat# 78500). Extension and all incubation procedures were performed using hybridization chamber and gaskets (Agilent #G2534A, #G2534-60003, #G2534-60011). Before binding reaction, microarrays blocked with 2% milk in PBS for 3-5 hours at room temp. Binding reactions carried out with ~0.25-4 μg of human HNF4α2 or HNF4α8 in NE from transfected COS-7 cells, or NE from WT and α7HMZ livers. NEs were diluted 1:10 in low salt PBM buffer (16 mM Hepes pH 7.8, 60 mM KCL, 8 mM EDTA, 8 mM EGTA) and processed through a 30 kDa cut-off column (Amicon, Cat# UFC503096) to a final concentration of 110 mM KCl and then applied to the arrays in PBM binding buffer (16 mM HEPES pH 7.8, 100 mM KCl, 8 mM EDTA, 8 mM EGTA, 0.1% Tween 20 plus 4-20 μg sonicated salmon sperm DNA). After 2 h of incubation, arrays were washed 3X for 3 min each with PBS plus 0.1% Tween 20. Mouse monoclonal anti-HNF4α P1/P2 (R&D Systems #PP-H1415-00) diluted 1:100 in PBS buffer plus 2% non-fat milk, 0.1% Tween 20 were applied directly to the slide and incubated for 24 h at RT, followed by a conjugated secondary Ab (GαM IgG [H+L] DyLight 550, Pierce Cat# 84540) diluted 1:50 (as described above) and then incubated for 90 min. Three washes, 3 min each, in PBS plus 0.1% Tween 20 were performed after each antibody incubation. HNF4α binding was imaged with 2-μm resolution using Agilent G2565CA Microarray Scanner at the UCLA DNA Microarray Core. Extraction and normalization of the data were as described previously (Bolotin et al., 2010). Position weight matrices (PWM) were generated using SeqLogo (Bembom 2017).

*Rapid Immunoprecipitation and Mass Spectrometry of Endogenous Proteins (RIME)*

**Hypotonic Buffer:** as in ChIP-seq

**Nuclei Lysis Buffer:** as in ChIP-seq

**IP Dilution Buffer:** as in ChIP-seq

**Shearing Buffer D3:** 0.1% SDS, 10 mM Tris-HCl pH 7.6, 1 mM EDTA in biology-grade water

**Inhibitors:** 2 μg/mL leupeptin, 2 μg/mL aprotinin, 1:100 protease inhibitor (Sigma #8340)

**RIPA buffer:** 15 mM Tris-HCl pH 8.0, 150 mM NaCl, 1% NP-40, 0.7% Deoxycholate

**DNaseI Buffer:** 40 mM Tris-HCl pH 8.0, 1 mM $CaCl_2$, 10 mM NaCl, 6 mM $MgCl_2$

RIME was performed as previously described (Mohammed et al., 2016) with the following modifications. WT and α7HMZ male mice (n=3, 16-18 weeks of age) were sacrificed at 10:30 (ZT3.5) and 20:30 (ZT13.5). Roughly 250-mg chunks of liver were collected from the same livers used for RNA-seq samples and fixed in 5 mL formaldehyde solution (1.1% MeOH-free formaldehyde, 2 μg/ml aprotinin, 2 μg/ml leupeptin in 1X PBS) for 10 min at RT. Cross-linking was stopped with 0.125 M glycine at RT for an additional 5 to 8 min. Samples were centrifuged (4 min at 2,500 rpm) at 4°C. The fixative was aspirated, the tissue immediately washed with cold PBS and centrifugation repeated. Fixed tissues were frozen and stored in liquid nitrogen.

Frozen tissues were placed in 1X PBS plus inhibitors and homogenized with a motorized homogenizer at 4°C as described above. The homogenized samples were passed through nylon cell strainer, dounced eight times, and centrifuged (5 min at 2,500 rpm) at 4°C. Cells were swelled in 1 mL Hypotonic Buffer for 10 min at 4°C, spun again to collect the nuclei, which were resuspended with Nuclei Lysis Buffer (see ChIP-seq for buffers), nutated for 20 min at 4°C and then gently resuspended in 0.5mL Shearing Buffer D3 (0.1% SDS, 10 mM Tris-HCl pH 7.6, 1 mM EDTA in biology-grade water) plus inhibitors. D3 buffer was added to fill the 1-ml sonication AFA milliTUBE (Covaris #520130). Samples were sonicated for 5.5 min, 30 s break, and again for 4 min in a Covaris S220 sonicator. Immediately after sonication samples were diluted 1:1 with IP dilution buffer plus all inhibitors. Samples were centrifuged (5 min at 11,000 rpm) for 5 min and the pellet, if any, was discarded. Before IP, samples were pre-cleared for 40 min at 4°C with 10 µL of pre-washed magnetic Protein A/G beads (Pierce #0088802). One day before sonication, magnetic Protein A/G beads (20 µL per sample) were pre-washed 3X with 1 mL PBS plus 0.05% Tween 20. For each sample, 3 µg of P1/P2 Ab or mouse IgG diluted in 300-400 µL PBS plus 0.05% Tween 20 and inhibitors were added to pre-washed beads and nutated for 20 h at 4°C. On the day of sonication, unbound Abs were removed from the beads, and the sonicated, precleared samples were added to the beads and nutated overnight at 4°C. The following day, the supernatant was removed and beads were washed 3X with 1 mL ice-cold RIPA buffer, and then 1X with 400 µL DNaseI Buffer and incubated in DNaseI buffer with 8 µL DNaseI enzyme (4 µg/µL, Sigma #D5319) for 20 min at 30°C. Afterwards, samples washed 3X with 0.5 mL RIPA buffer at room temperature and then 2X with 1 mL ice-cold RIPA buffer. Parallel IP samples (± DNA digestion) were examined for the presence of DNA by Qubit fluorometer,  to confirm high efficiency digestion. IP'd material was washed 2x with 1 mL 50 mM $NH_4CO_3$. At the last wash, the suspension was transferred to a new non-stick tube. The wash buffer was removed and the IP beads were immediately frozen and later subjected to mass spectrometry as described below.

Multidimensional protein identification technology (MudPIT) analysis was performed by the Proteomics Core Facility in the IIGB at the University of California, Riverside. Sample preparation following IP analysed by 2D nano-liquid chromatography tandem MS (2D nano-LC/MS/MS). Briefly, following IP, beads were washed in trypsin buffer [50 mM ammonium bicarbonate, 10% (vol/vol) acetonitrile] and digested overnight at 37 °C (1μg trypsin in 100 μL buffer) and washed one time [10 min 100 μL 50% (vol/vol) acetonitrile, 5% (vol/vol) acetic acid]. Digest supernatant and post-digest wash supernatant were combined. Tryptic peptides were pelleted by SpeedVac concentrator and redissolved in 20 μL 0.1% formic acid. Peptides were separated by a 2D nano-Acquity ultra-performance LC system (2D nano-UPLC) (Waters) and analyzed with an Orbitrap Fusion MS system (Thermo Fisher). High-pH reversed-phase LC with 20 mM ammonium formate pH 10 (solvent A) and 100% (vol/vol) acetonitrile (solvent B) was used to fractionate the tryptic peptides into five fractions on an XBridge BEH130 C18 trap column [5 μm particle, 300-μm internal diameter (i.d.), 5-cm long; Waters #186003682]. Five fractions and a flush fraction were collected at (1) 13%, (2) 18%, (3) 21.5%, (4), 27%, (5) 50%, and a final flush of 60% solvent B. Each and every of these fractions was first concentrated and then separated with a conventional reverse phase gradient in acidic condition. A Symmetry C18 column (5-µm particle, 180-µm i.d., 20-mm long, Waters #186003514) was used to concentrate and desalt the peptides of each fraction. The samples were further separated on a BEH130 C18 column (1.7-µm particle,75-µm i.d., 20-cm long, Waters #186003544). The mobile phase A and B solvents for separation gradient were 0.2% formic acid in water and 0.2% formic acid in acetonitrile, respectively. The mobile phase nano-flow rate was 0.3 µL/min with the following 1-hour gradient: 0–1 min, 3%B; 1–30 min, 50% B; 30–31 min, 85%B; 31–35 min, 85% B; 35–36 min, 1% B; and 36–60 min, 1% B.

The MS analysis part of MudPIT was carried out with Orbitrap Fusion MS system (Thermo Fisher). A data-dependent acquisition (DDA) survey method using HCD (high-energy collision dissociation) fragmentation technique was employed in a positive ion mode. The instrument parameter included ESI spray voltage at 2300 V, ion transfer tube temperature 275°C, and 0 sweep gas. MS1 scan was carried out with Orbitrap mass analyzer with its resolution set at 120,000 and normal mass range from 300 to 2000 m/z. S-Lens RF level was 60%. AGC target was set 200,000. 50 msec was set for maximal injection time. Microscan was set for 1. For MS2 scan, top-speed scanning method was used with time window of 4 seconds. Monoisotopic selection was allowed, peptide ions with charge state from 2 to 5 and other undetermined charge states were all selected for MS2 fragmentation by HCD. Dynamic exclusion was activated after three MS2 spectra were acquired for each m/z within 1 min window. Dynamic exclusion duration was for 5 min, and exclusion mass window was +/- 30 ppm. Ion selection for MS2 acquisition was arranged from most intense peak to least intense peak with minimal intensity of NL 100,000. For HCD fragmentation, quadrupole isolation window was fixed at 2 m/z, collision energy was set at 30%, ion trap was chosen as mass detector to collect all MS2 fragments for each individual peptide ions. Ion trap scan rate was set at rapid with scan range set at normal. First fragment mass was set at 120 m/z. AGC target was set at 10,000 with maximal injection time of 0.1 second. All raw MS1 and MS2 spectra were processed with Proteome Discoverer 2.1 (Thermo Scientific) to generate mgf files, which were then submitted to Mascot searching engine to match against NCBI non-redundant mouse protein database for protein identification. Only proteins with 1% FDR cut-off ($q \leq 0.01$) were considered for subsequent analysis.

Area under the curve, as reported by Proteome Discoverer, were averaged together for WT and α7HMZ samples (n=3). IgG samples (n=3) from both WT and α7HMZ at 10:30 AM were averaged together to create a background sample for the 10:30 analysis. There was one pooled IgG sample at 20:30, which was combined with the pooled 10:30 AM Ig background for the 20:30 analysis. Areas were converted to log2 scale and the fold-change above IgG background was calculated for the WT and α7HMZ samples. Proteins with less than 8-fold change above background were omitted. Similarly, a 8-fold difference between WT and α7HMZ samples was used to identify unique protein interactions.

*Primary metabolite and complex lipids analysis*

All metabolomic analysis was performed at the West Coast Metabolomics Center at the University of California Davis as described before (Deol et al., 2017). Primary metabolites and complex lipid analysis was done on livers from WT (C57BL/6N) and α7HMZ (backcrossed 10+ generations into C67BL/6N) adult male mice harvested mid-morning and fed the standard chow (n=8, aged 38 weeks). Liver tissue was rinsed in cold PBS, snap frozen and stored in liquid nitrogen until analysis. Concentrations are presented as pmol/gm in tissue. Fold-enrichment was performed using MetaboAnalyst. One outlier from each group (a7HMZ.4, WT.4) was removed before plotting and statistical analysis. Data are presented as mean +/- standard error of mean (SEM). Student's T-test was used to determine statistical significance ($p < 0.05$) using GraphPad Prism v6.

*Oxylipin analysis*

Analysis of non-esterified oxylipins, was performed on a mixed 129/Sv plus C57BL/6 background (n=3 per group, aged 12-13 weeks) (Briançon and Weiss, 2006). Liver tissue was rinsed in cold PBS, snap frozen and stored in liquid nitrogen until analysis. Tissue homogenates (100 mg) were extracted and analyzed according to previously described protocols (Matyash et al., 2008; Yang et al., 2009). Briefly, samples were extracted by solid phase extraction and analyzed by ultrahigh performance liquid chromatography tandem mass spectrometry (UPLC-MS/MS) (Agilent 1200SL-AB Sciex 4000 QTrap). Analyst software v.1.4.2 was used to quantify peaks according to corresponding standard curves with their corresponding internal standards.

Oxylipin concentrations are presented as pmol/gm in tissue. Data are presented as mean +/- standard error of mean (SEM). Student's T-test was used to determine statistical significance ($p < 0.05$) using GraphPad Prism v6.

**QUANTIFICATION AND STATISTICAL ANALYSIS**

Differential gene expression (DEG) was measured using raw read counts with DESeq2: statistical significance was defined as adjusted p-value (padj.) ≤ 0.01, unless otherwise noted. Legends denote any thresholds using log2 fold change (log2FC) cutoffs. R library "gage" was utilized to identify differentially enriched KEGG pathways. ChIP-seq peaks were called with MACS2 and then filtered on -log10(P-value) ≥ 10, i.e., p-value ≤ 1e-10, to approach six-fold enrichment above control. Differentially bound peaks were identified using DiffBind with MACS2 output. RIME samples were analyzed with Proteome Discoverer 2.0: areas reported were converted to a log2 scale, thus fold changes were calculated on the log2 scale. Methods to filter RIME data discussed above. All heatmaps were generated with pheatmap package in R. Heatmap data were row-normalized before plotting with the exception of NR heatmap in Figure 2. All barplots represent mean ± SEM. Transcription Factor (TF) rankings for Cleveland plots were ordered at the 13:30 (peak HNF4α expression) then manually curated with the aid of PANTHER; all TF genes with FPKM > 50 plotted using the 10:30 FPKM values. All Venn diagrams were generated with VennDiagram package in R. Unique and common RIME results were submitted to DAVID for ontology analysis. Statistical significance for metabolite data defined as padj. ≤ 0.05 by Mann-Whitney U-test. All barplots represent mean ± SEM; significant differences are noted between genotypes at a given time point, unless indicated otherwise. For FPKM plots, padj values are from DESeq2; in other plots, p-values are from two-way Student's T-test or One/Two-way ANOVA, as indicated.

*External expression datasets*

The following mouse differential expression analyses were used to generate scatterplots for Figures 1GH and S1AD. The HNF4α mouse liver knockout (KO) data was generated by (Walesky et al., 2013) using Affymetrix Mouse430_2.0 genechips. Data were summarized to the gene level taking the largest fold change value and associated p-value for a single gene if more than one transcript was reported. The adult vs. fetal differential expression analysis was generated by the ENCODE project (ENCSR000BYS, ENCSR000BZI) (ENCODE Project Consortium, 2012). Gene quantifications files were downloaded and RSEM expected_count values treated as raw read counts for differential expression analysis with DESeq2, with log2 Fold Change (log2FC) and adjusted p-values (padj.) used for plotting and highlighting values in scatterplots. The C57BL/6 vs Hepa1-6 and Hepa1c1c7 differential expression analysis was generated with DESeq2 (Rudolph et al., 2016). Similarly, log2FC and padj were used for plotting and highlighting values in scatter plots, which were graphed using ggplot2 library in R. All scatterplots used the RNA-seq from WT vs. α7HMZ adult male livers at 10:30 AM (fed) described in this study.

**DATA AND SOFTWARE AVAILABILITY**

The raw and processed RNA-seq data have been deposited in GEO under GSE117972.

The raw and processed ChIP-seq data have been deposited in GEO under _in progress___.

The processed PBM data have been deposited in the Nuclear Receptor DNA Binding Project (NRDBP) under _in progress___.

The raw metabolomics data (primary metabolites and complex lipids) have been deposited in Metabolomics Workbench (www.metabolomicsworkbench.com) under Project #PR000461.

### ADDITIONAL RESOURCES

Nuclear Receptor DNA Binding Project: http://fmslab.ucr.edu/binplone

The NRDBP is a project dedicated to the understanding of DNA binding preferences of nuclear receptors. This website serves as a centralized resource for protein binding microarray (PBM) data related to this project, and hosts the PBM results utilized in this experiment.
